# BRCA2 loss drives a PKM1-dominant glycolytic state that is selectively lethal to pyruvate kinase activation

**DOI:** 10.64898/2026.09.29.754041

**Authors:** Maira Di Tano, Mujmmail Ahmed, Shiri Li, Michael Lyashenko, Neil Ruthen, Ethan Tse, Isaac Nathoo, Carolina E. Echeverria, Jack Sanford, Anas Saleh, Jeshua Kim, Ezequiel Dantas, Davide Pradella, Kyu Y. Rhee, Ed Reznik, Lewis C. Cantley, Marcus D. Goncalves

**Affiliations:** Department of Medicine, New York University Grossman School of Medicine, NYU Langone Health, New York, NY, USA 10016; Department of Medicine, Weill Cornell Medicine, New York, NY, USA 10021; Memorial Sloan Kettering Cancer Center, New York, NY, USA 10065; Weill Cornell Graduate School of Medical Sciences, Weill Cornell Medicine, New York, NY 10021, USA; Weill Cornell-Rockefeller-Sloan Kettering Tri-Institutional MD-PhD Program, New York, NY, US; Dana-Farber Cancer Institute, Boston, MA, USA 02215

## Abstract

Homologous recombination deficiency (HRD), most commonly arising from BRCA1/2 loss, is accompanied by metabolic remodeling whose contours and therapeutic implications remain poorly defined. Through pan-cancer analysis of 10,619 tumors from The Cancer Genome Atlas (TCGA) and isogenic BRCA1/2-deficient models, we show that HRD tumors display broad metabolic rewiring, with consistent alterations in glucose metabolism. Unexpectedly, BRCA1/2 loss is associated with selective upregulation of the constitutively active glycolytic enzyme Pyruvate Kinase M1 (PKM1) isoform, contrasting with the PKM2-dominant state of most cancers, and produces an elevated pyruvate kinase activity state. Overactivating PKM, with small molecules activators, including the FDA-approved drug mitapivat, triggers superoxide accumulation, disrupts PINK1-dependent mitochondrial quality control, induces DNA damage, and selectively kills BRCA2-deficient tumors *in vivo*, while sparing BRCA2-proficient counterparts. These findings recast HRD as a metabolic state and identify PKM activation as a candidate therapeutic strategy for biomarker-selected HR-deficient cancers.

**Significance:** BRCA1/2-deficient cancers rely on an unusual PKM1-dominant glycolytic state. We show that pushing activity past a critical threshold by pharmacologically overactivating PKM triggers oxidative stress, disrupts mitochondrial quality control, and selectively kills BRCA2-deficient tumors. This reveals a druggable metabolic vulnerability in HRD cancers and repositions existing therapy for biomarker-guided treatment.

## Introduction

Genome maintenance and cellular metabolism are intimately coupled. DNA repair pathways consume deoxyribonucleotide triphosphates (dNTPs), nicotinamide adenine dinucleotide (NAD+), S-adenosylmethionine and adenosine triphosphate (ATP), while metabolic state shapes the fidelity and efficiency of repair (1–3). This bidirectional relationship implies that cells with chronically elevated DNA damage or compromised repair operate under metabolic constraints that differ from those of their repair-competent counterparts. Cancers harboring defects in homologous recombination (HR) are a paradigmatic example. HR-deficient (HRD) tumors sustain ongoing replication stress that is expected to impose persistent metabolic demand. Consistent with this view, HRD has been linked to distinct redox and oxidative-phosphorylation states (4). Yet a systematic view of how HRD reshapes central metabolism, and whether this reshaping itself creates therapeutic liabilities, remains elusive.

Among inherited cancer predispositions, germline mutations in *BRCA1* and *BRCA2* are the most common cause of HRD and confer markedly elevated risk of breast, ovarian, prostate and pancreatic cancer (5–7). BRCA1 and BRCA2 act at distinct but converging steps of HR. BRCA1 acts in end resection and pathway choice, while BRCA2 supports RAD51 loading onto resected DNA fragments. Their loss commits cells to error-prone repair through non-homologous end joining and microhomology-mediated alternatives, generating the genomic scars that define the HRD phenotype (8). Whether the metabolic state imposed by HR loss provides an orthogonal axis of vulnerability has become a question of both mechanistic and translational importance.

Among the metabolic axes most consistently rewired in cancer is glucose handling. Most tumor cells favor aerobic glycolysis, the fermentation of glucose to lactate irrespective of oxygen availability, over complete pyruvate oxidation, a phenomenon known as the Warburg effect (9,10). Recent works indicate that this preference arises in large part from the constraints of NAD+ homeostasis: cells engage aerobic glycolysis when mitochondrial NAD^+^ regeneration cannot keep pace with cellular demand, either because oxidation reactions outstrip ATP turnover or because glycolytic flux exceeds the capacity of the malate-aspartate and glycerol-3-phosphate shuttles to deliver reducing equivalents to the mitochondria (11,12). The partitioning of glucose carbon between fermentation and oxidation is set largely at the final step of glycolysis, controlled by pyruvate kinase M (PKM), which catalyzes the conversion of phosphoenolpyruvate (PEP) and ADP to pyruvate and ATP. PKM activity is therefore a central node coupling glycolytic throughput to mitochondrial oxidative capacity, and its dysregulation can shift cells between metabolic states with very different demands on NAD+ regeneration.

Alternative splicing of *PKM*, via mutually exclusive inclusion of exon 9 or exon 10, generates two splicing isoforms, PKM1 and PKM2 (13,14). PKM1 is constitutively active, whereas PKM2 exists in equilibrium between low- and high-activity states governed by allosteric metabolites and post-translational modifications (15). Most proliferating tissues, including most cancers, predominantly express PKM2. Its regulatory flexibility is thought to enable rapid switching between catabolic and anabolic glucose handling as cellular demands shift, therefore promoting proliferation (13–16). This isoform difference has motivated the development of small-molecule PKM2 activators (including DASA-58, TEPP-46, and the clinically approved drug mitapivat) that lock PKM2 into a high-activity conformation resembling PKM1, thereby restricting glucose flux into biosynthetic side branches and suppressing tumorigenesis (17–19). Despite the availability of these compounds, the clinical development of PK activators in oncology has been limited by uncertainty over which tumors might respond.

Here, we use pan-cancer analysis of The Cancer Genome Atlas (TCGA) tumors and isogenic BRCA1/2-deficient cell models to define the metabolic state imposed by HRD. We show that loss of BRCA1/2 alters glycolysis by upregulating PKM1, which enhances the sensitivity to PK activators.

## Results

### PKM expression and activity are elevated in BRCA2-deficient tumors through upregulation of the PKM1 isoform *in vitro* and *in vivo*

To determine whether HRD tumors exhibit metabolic features distinct from HR-proficient tumors, we performed a pan-cancer analysis of 10,619 tumors from TCGA (20) stratified according to their HRD score (21). Increasing HRD score was associated with broad metabolic rewiring across tumors (Fig.1a, Supplementary Fig.1A). Among the pathways most consistently altered in HRD tumors were glucose metabolism, including glycolysis/gluconeogenesis and oxidative phosphorylation (Fig.1A, Supplementary Fig.1A). These metabolic alterations were independently validated at the protein level using data from the Clinical Proteomic Tumor Analysis Consortium (CPTAC) (Supplementary Fig.S2A) (22).

**Figure 1.**
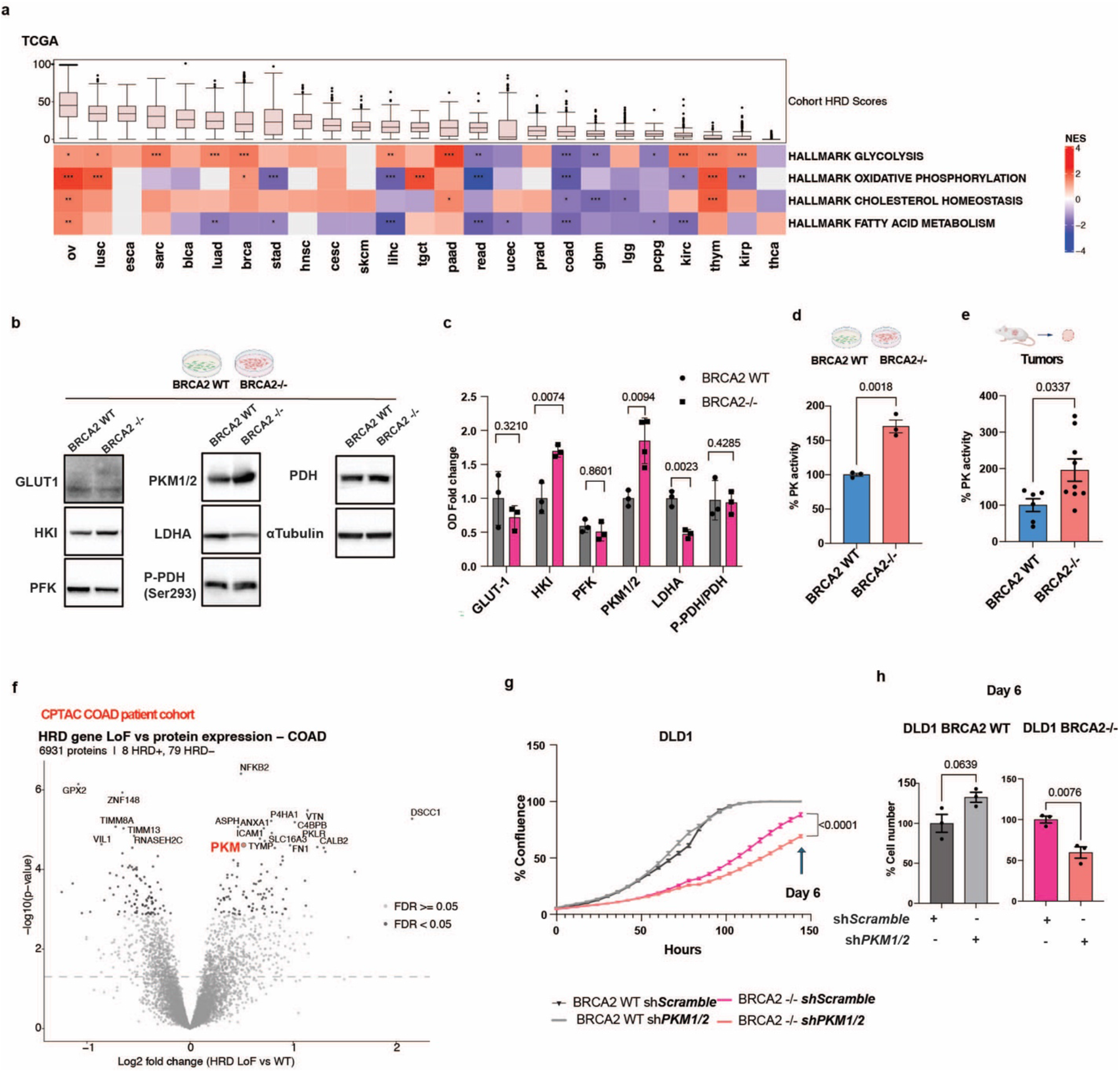
PKM activity is upregulated in BRCA2-deficient tumors and sustain cell growth. **a** Heatmap showing normalized enrichment scores (NES) for significantly enriched Hallmark gene sets identified performed across 10,619 TCGA cancer types stratified by HRD score. Cell color denotes the NES, with red indicating positive enrichment and blue indicating negative enrichment. Positively enriched pathways (red) are progressively upregulated with higher HRD score, whereas negatively enriched pathways (blue) are progressively downregulated. Color intensity reflects the relative magnitude of regulation. Metabolism-related pathways are shown in tumors with sample size≥ 100. Multiple testing correction was applied using the Benjamini–Hochberg method, **b** Immunoblot analysis of GLUT1, HKI, PFK, PKM, LDHA, phospho-PDH, and PDH in DLD1 BRCA2 WT and BRCA2-/- cells. α-Tubulin was used as a loading control. A representative image is shown (n=3). **c** Quantification of immunoblots shown in **b** (n=3, PKM1/2 in BRCA2-/- n=4). P value was calculated using the two-sided unpaired Student’s *t*-tests. **d** Pyruvate kinase (PK) activity in DLD1 cells grown *in vitro* (n=3). **e** PK activity assay in DLD1 mouse-derived tumor masses (DLD1 BRCA2 WT n=6, DLD1 BRCA2-/- n=9). Schematic of cells and mouse tumors were created with Biorender. P value was calculated using the two-sided unpaired Student’s *t*-tests. **f** Volcano plot showing differential protein expression in the CPTAC colorectal adenocarcinoma (COAD) patient cohort (HR deficient n=8, HR proficient n=79), with PKM highlighted in red. **g** Growth curve of DLD1 BRCA2 WT and BRCA2-/- infected with sh*Scramble* or sh*PKM1/2*. Percentage of confluence over time is shown (n=8). **h** Cell number 6 days (144 hours) after seeding is shown (n=3). P value was calculated using the two-sided unpaired Student’s *t*-tests. All data are presented as mean ± S.E.M. P values were determined by two-sided unpaired t-test.

To explore the effects of HRD on these pathways, we measured transcriptional and protein changes in isogenic DLD1 colorectal cancer cells with (WT) and without BRCA2 expression (BRCA2-/-), a well-established and widely used model of BRCA2-mediated HRD. Consistent with the TCGA data, glycolysis was among the most significantly regulated transcriptional pathways (Supplementary Fig.S3A). We observed protein-level changes in key glycolytic enzymes, including increased HK1 and PKM, and reduced LDHA (Fig.1B-C). Because PKM determines glucose fate, we confirmed that pyruvate kinase activity was higher in BRCA2-deficient cells in culture and in tumors in mice (Fig.1D-E). CPTAC colorectal adenocarcinoma data (COAD) likewise identified PKM as one of the proteins significantly upregulated in HRD tumors relative to HR-proficient tumors (Fig.1F).

Next, we investigated why BRCA2-deficient cells upregulate PKM and whether this is a pro-survival mechanism. We depleted PKM in both BRCA2 WT and BRCA2-/- DLD1 cells (Supplementary Fig.S4A). PKM knockdown had no detectable effect on growth of BRCA2 WT cells; in contrast, it significantly impaired growth of BRCA2-/- cells, measured by cell confluence and total cell number (Fig.1G-H), supporting the idea that BRCA2-deficient cells are selectively dependent on PKM for proliferation.

Because PKM exists as alternatively spliced isoforms, PKM1 and PKM2, we asked whether increased PKM activity was associated with altered isoform expression. We found selective upregulation of PKM1 at the mRNA, protein, and activity levels upon BRCA2 loss (Fig.2A-D). However, RNA-seq analysis did not reveal significant transcriptional changes in established PKM splicing regulators, including PTBP1, PTBP2, hnRNPA1, hnRNPA2/B1, SRSF3, RBM4, and KHDRBS1, pointing to other post-transcriptional or post-translational mechanisms (Supplementary Fig.S5A).

**Figure 2.**
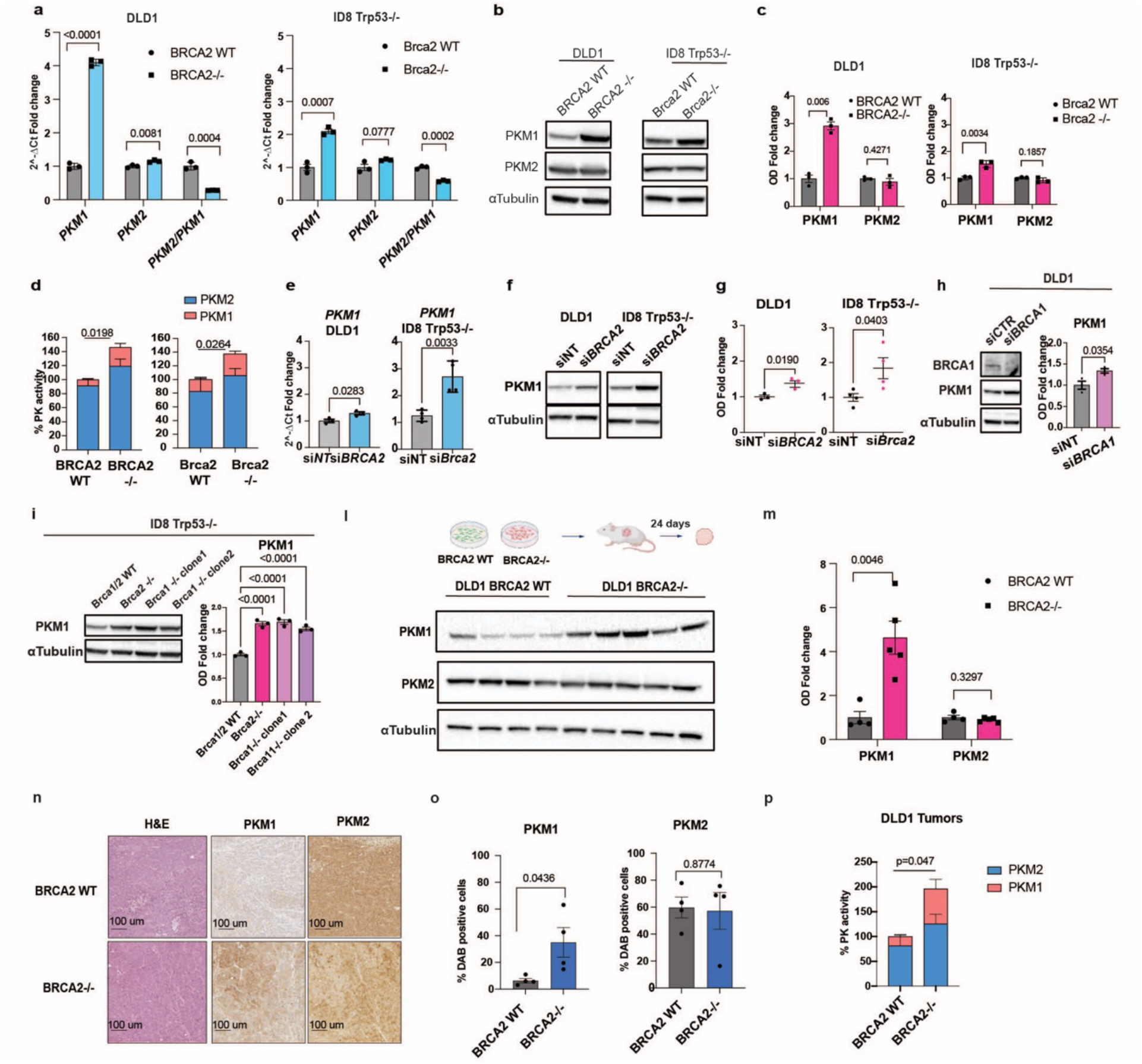
BRCA1/2 loss drives selective PKM1 isoform upregulation at the transcript, protein, and activity levels. **a** *PKM1* and *PKM2* mRNA levels in DLD1 BRCA2 WT and BRCA2-/- cells (left) and ID8 Trp53-/- Brca2 WT and Brca2-/- cells (right), measured by qPCR (n=3). **b** Immunoblot analysis of PKM1 and PKM2 in the indicated cell lines. α-Tubulin was used as a loading control. A representative image is shown (n=3). **c** Quantification of immunoblots shown in **b** (n=3). **d** PK activity assay in DLD1 (n=4) and ID8 Trp53-/- (n=3). **e** *PKM1* mRNA levels in DLD1 cells (n=3) and ID8 Trp53-/- (n=4) transfected with non-targeting siRNA (siNT) or siRNA directed against BRCA2. **f** Immunoblot analysis of PKM1 under the indicated conditions in DLD1 cells (left, n=3) and ID8 Trp53-/- cells (right, n=4). α-Tubulin was used as a loading control. A representative image is shown. **g** Quantification of immunoblots shown in **f**. **h** Immunoblot analysis of BRCA1 and PKM1 in DLD1 cells transfected with si*NT* or si*BRCA1* (left), with quantification shown on the right. A representative image is shown (n=3). **i** Immunoblot analysis of PKM1 in ID8 Trp53-/- Brca1/2 WT, Brca2-/-, and two independent Brca1-/- clones (cl26.1 and cl36.1) (left), with quantification shown on the right (n=3). **l** DLD1 BRCA2 WT and BRCA2-/- cells were implanted into mice and tumors were collected after 24 days (schematic, top). Immunoblot analysis of PKM1 and PKM2 in tumors derived from DLD1 BRCA2 WT (n=4) and BRCA2-/- cells (n=5). α-Tubulin was used as a loading control. A representative image is shown. Quantification of immunoblots shown in **l**. **n** Immunohistochemistry (IHC) of mouse-derived DLD1 tumor sections stained for PKM1 and PKM2 (n=4). H&E staining is shown on the left. Scale bar 100μm. **o** Quantification of PKM1 and PKM2 IHC staining (n=4). **p** PK activity assay in mouse-derived DLD1 tumors (DLD1 BRCA2 WT n=6, DLD1 BRCA2-/- n=9). Data are presented as mean ± S.E.M. P values were determined by two-sided unpaired t-test

We confirmed these findings in an independent HRD model using murine ID8 cells lacking p53 (ID8 Trp53-/-) with and without BRCA2 (23–25) (Fig. 2A-D). To exclude long-term adaptation of selected knockout clones, we performed acute knockdown of BRCA2/Brca2 in DLD1 and ID8 Trp53-/- cells (Supplementary Fig.S5B-D). After 48 hours, PKM1 mRNA and protein were significantly increased (Fig.2E-G). Similarly, PKM1 was acutely upregulated after *Brca2* knockdown in the murine pancreatic cancer cell line K8082, suggesting a conserved metabolic response to HRD across colorectal, ovarian, and pancreatic cancer models (Supplementary Fig. S5E-H) (26). Next, we explored whether PKM1 upregulation was specific to BRCA2 loss or a more general response to HRD. We performed BRCA1 knockdown in DLD1 cells and used two published ID8 clones with CRISPR-mediated BRCA1 deletion (25). In each case, we observed increased PKM1 protein expression (Fig.2H-I), suggesting that this metabolic change may be a general response to HRD.

We then asked whether the PKM1-selective overexpression observed in cultured cells extends to tumors. WT and BRCA2-/- DLD1 cells were implanted subcutaneously into nude mice, and tumors were collected after 24 days. In agreement with our *in vitro* data, PKM1 expression and activity were selectively increased in BRCA2-/- tumors, whereas PKM2 levels were not significantly altered (Fig.2L-P). Together, these data suggest that PKM expression and activity are elevated in BRCA1/2-deficient tumors through upregulation of the PKM1 isoform, and that this increase in PKM activity is required to sustain tumor growth.

### BRCA2-deficient tumor cells are hypersensitive to PKM activation, and PKM1 overexpression is sufficient to confer this vulnerability

Pharmacological PK activators stabilize PKM2 in a high-activity tetrameric state and have been shown to compromise cellular fitness in certain settings (17,27). We reasoned that BRCA2-deficient cells, which already exhibit elevated PK activity through PKM1 upregulation, might respond differently to further pharmacological activation than HR-proficient cells. We therefore asked whether the altered PKM state in BRCA2-deficient cells created a targetable vulnerability. Treatment of DLD1 and ID8 *Trp53-/-* cells with the PK activator, DASA-58, exerted a cytostatic effect in both BRCA2-proficient and BRCA2-deficient backgrounds (Supplementary Fig.S6). However, DASA-58 selectively induced cell death in BRCA2-deficient cells, while sparing BRCA2 WT counterparts (Fig. 3A). This selective toxicity was accompanied by increased DNA damage, as indicated by increased histone H2AX phosphorylation (Fig. 3B).

**Figure 3.**
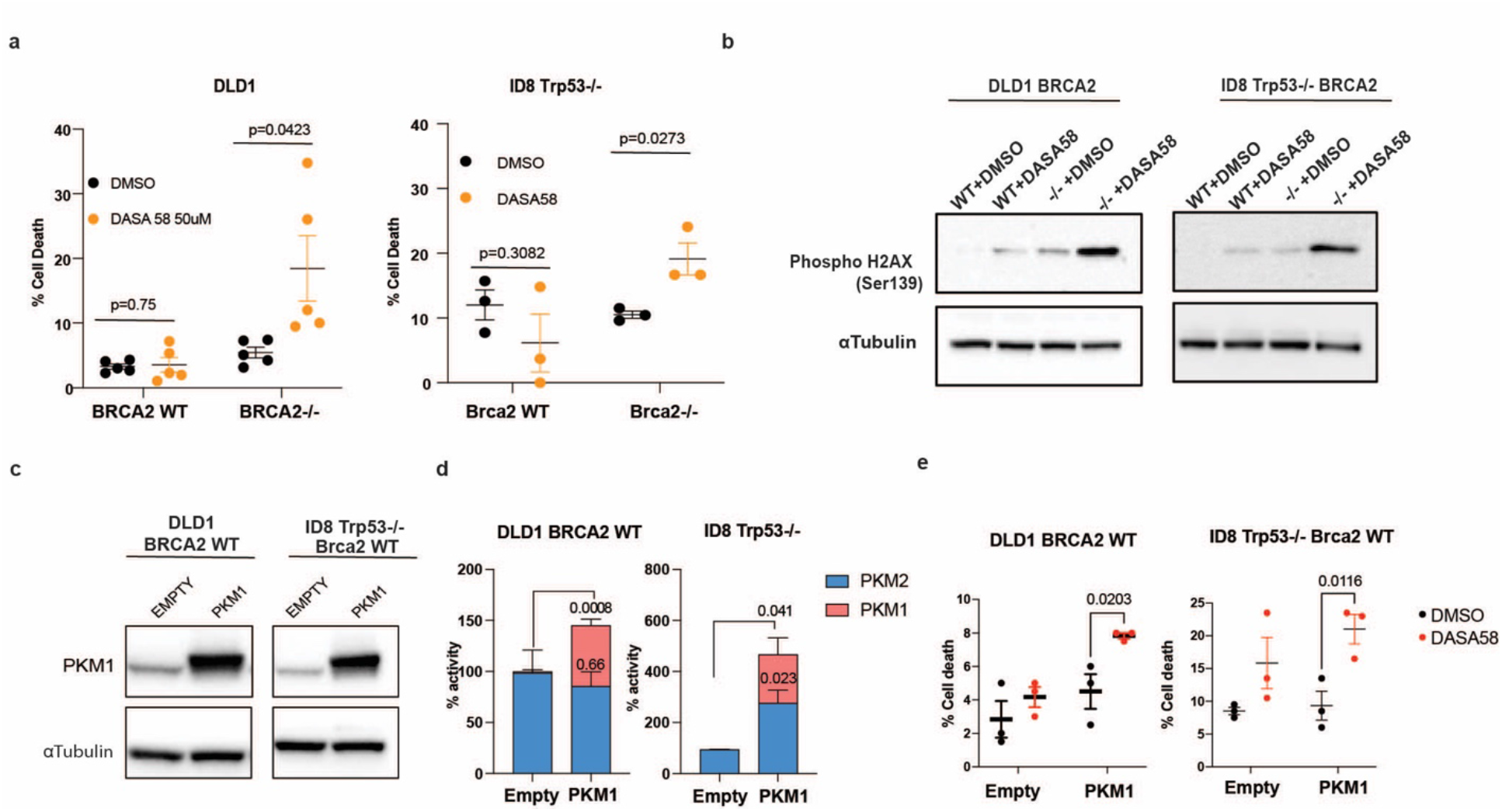
PKM1 overexpression is sufficient to confer sensitivity to PKM activation in BRCA2-proficient cancer cells. **a** Cell death measured by Trypan blue assay in DLD1 BRCA2 WT and BRCA2-/- cells (left, n = 5) and ID8 Trp53-/- Brca2 WT and Brca2-/- cells (right, n = 3) treated with DASA58 (50 μM) or DMSO. **b** Immunoblot analysis of phospho-H2AX under the indicated conditions. A representative image is shown (n=3). α-Tubulin was used as a loading control. **c** Immunoblot analysis of PKM1 in DLD1 and ID8 Trp53-/- BRCA2/Brca2 WT cells transfected with empty vector or stably overexpressing PKM1. A representative image is shown (n=3). α-Tubulin was used as a loading control. A representative image is shown. **d** PK activity assay on DLD1 BRCA2 WT and BRCA2-/- (n = 3). **e** Cell death measured by Erythrosine B assay in DLD1 and ID8 Trp53-/- BRCA2/Brca2 WT cells transfected with empty vector or stably overexpressing PKM1 and treated with DASA58 (50 μM) or DMSO (n = 3). Data are presented as mean ± S.E.M. P values were determined by two-sided unpaired t-test.

To determine whether elevated PK activity is sufficient for this response, we overexpressed PKM1 in BRCA2 WT cells (Fig. 3C-D). PKM1 overexpression was sufficient to sensitize BRCA2-proficient cells to DASA-58, as shown by increased cell death (Fig. 3E), indicating that the PKM1-high state is a functional determinant of this drug’s response.

### Mitochondrial oxidative stress drives cell death in BRCA2-deficient cells upon PK activation

We next asked whether the changes in glucose handling and PK activity in BRCA2-deficient cells were associated with altered mitochondrial function. BRCA2-deficient DLD1 cells showed a significant reduction in basal oxygen consumption rate (OCR) relative to WT cells (Fig. 4A). While there was no change in mitochondrial number by mtDNA counts or abundance of the mitochondrial proteins TOM20 or TIM23 (Supplementary Fig. S7A-B), immunoblotting with an OXPHOS antibody cocktail revealed decreased abundance of respiratory chain complexes I, II, and IV (Fig. 4B). These results suggest a specific defect in oxidative capacity rather than a global change in mitochondrial mass.

**Figure 4.**
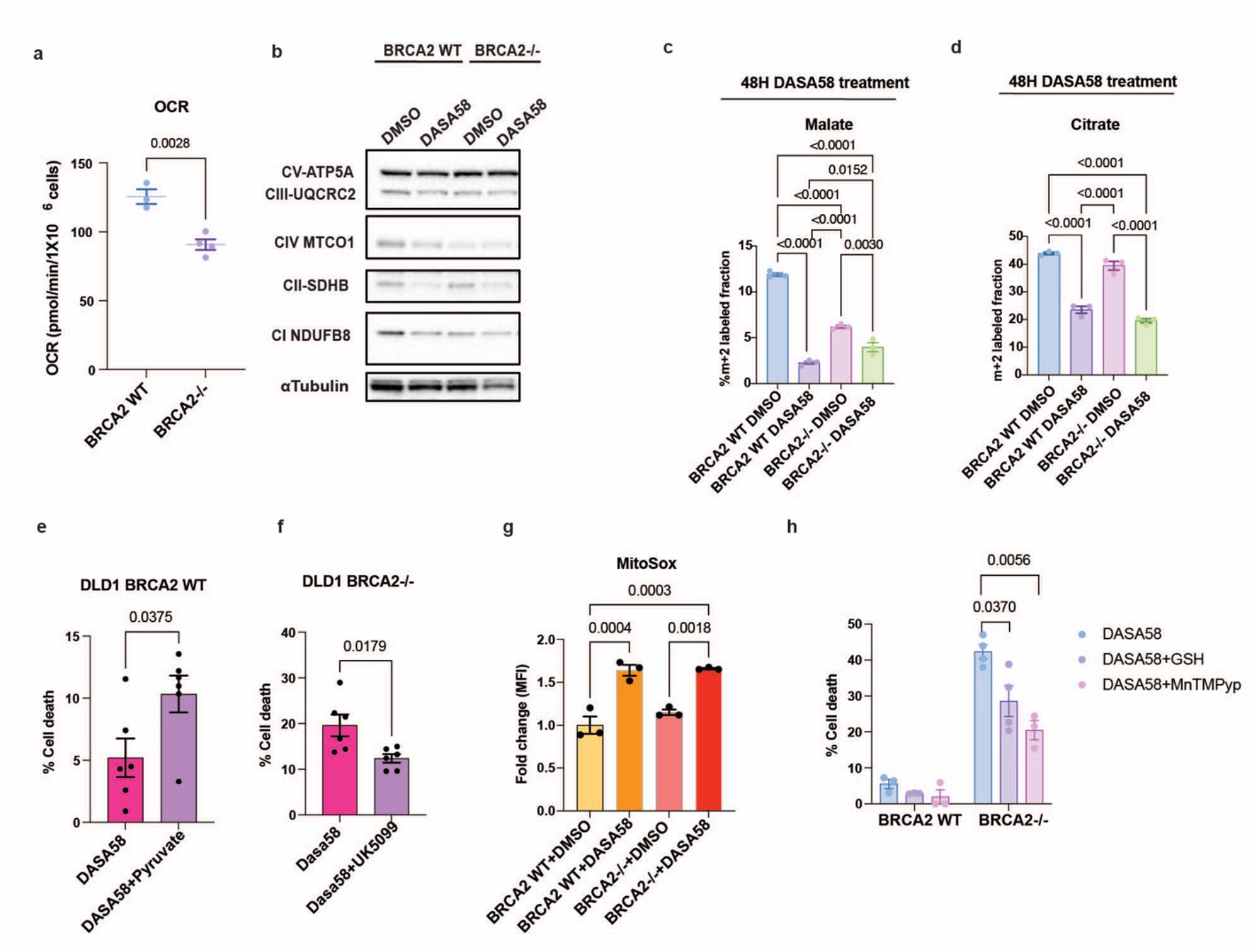
Pyruvate kinase activation compromises mitochondrial function and drives ROS-dependent cell death in BRCA2-deficient cells. **a** Oxygen consumption rate (OCR) measured by Seahorse analyzer in DLD1 BRCA2 WT (n=3) and BRCA2-/- cells (n=4). b Immunoblot analysis of respiratory chain complexes in DLD1 BRCA2 WT and BRCA2-/- cells treated with DASA58 or vehicle. A representative image is shown (n=3). α-Tubulin was used as a loading control. A representative image is shown. c-d M+3-labeled (^13^C incorporation) malate (2h labelling) (c) and citrate (4h labelling) (d) in DLD1 cells (n = 3). e Cell death measured by Erythrosine B assay in DLD1 BRCA2 WT cells supplemented with or without pyruvate and treated with DASA58 (n = 6). **f** Cell death measured by Erythrosine B assay in DLD1 BRCA2-/- cells treated with or without the mitochondrial pyruvate carrier inhibitor UK5099 and DASA58 (n = 6). **g** Mitochondrial superoxide levels measured by MitoSOX Red and flow cytometry (n = 3). **h** Cell death measured by Erythrosine B assay in DLD1 BRCA2 WT (n=3) and BRCA2-/- cells treated with DASA58, with or without glutathione (GSH) (n=4) or MnTMPyP (n = 3). Data are presented as mean ± S.E.M. P values were determined by one-way ANOVA.

To assess glucose oxidation directly, we performed steady-state isotopic labeling with [U-^13^C]-glucose. BRCA2-/- cells incorporated less [^13^C] label into the tricarboxylic acid (TCA) cycle intermediates, citrate and malate (Fig. 4C-D), indicating diminished entry of glucose-derived carbon into the TCA cycle. Despite the reduced [^13^C] incorporation, the absolute pool sizes of TCA intermediates were increased in BRCA2-/- cells (Supplementary Fig. S8A), suggestive of alternative anaplerotic substrates feeding the TCA cycle.

To examine the metabolic consequences of PK activation, we extended our metabolomic analysis in DLD1 treated with DASA-58. As expected, DASA-58 increased the pyruvate to PEP ratio in both WT and BRCA2-/- cells, confirming engagement of pyruvate kinase (Supplementary Fig. S8B). Acute exposure (1h) to DASA58 treatment led to a reduction of TCA cycle intermediates’ total abundance, including malate and fumarate, in both genotypes (Supplementary Fig. S8A). Intracellular lactate and alanine abundance increases selectively in BRCA2-/- cells treated with the PK activator (Supplementary Fig.S8C, Supplementary Table 1). These findings suggest that pharmacologic PK activation in BRCA2-/- cells generates pyruvate excess and perturbs its downstream partitioning, leading to accumulation of pyruvate-derived metabolites (lactate and alanine) while reducing TCA cycle metabolite pools.

We next assessed the effects of prolonged PK activation. After 48 hours of DASA-58 treatment, [U-^13^C]-glucose tracing revealed sustained reductions in [^13^C] incorporation into TCA cycle intermediates, consistent with durable suppression of glucose-derived carbon entry into the TCA cycle (Fig. 4C-D). Lactate labeling was also reduced, indicating that the transient rise in intracellular lactate abundance did not reflect sustained enhancement of glucose-derived lactate production (Supplementary Fig. S8D, Supplementary Table 2). Instead, these findings suggest that lactate production constitutes only a transient and partial outlet for excess pyruvate.

We therefore asked whether the reduction in TCA cycle intermediates reflected impaired mitochondrial pyruvate utilization rather than reduced pyruvate availability per se. If so, increasing pyruvate availability should sensitize BRCA2-proficient cells to PK activation, whereas blocking mitochondrial pyruvate import should protect BRCA2-deficient cells. Consistent with this model, exogenous pyruvate sensitized WT cells to DASA-58 (Fig. 4E). Conversely, inhibition of the mitochondrial pyruvate carrier with UK5099 rescued DASA-58-induced cell death in BRCA2-deficient cells (Fig. 4F). Together, these data indicate that mitochondrial pyruvate entry remains the proximal driver of selective cytotoxicity in BRCA2-deficient tumor cells even though some excess pyruvate is diverted into lactate and alanine.

Excess mitochondrial pyruvate is known to drive electron flux through the respiratory chain and, when this flux exceeds the capacity of the chain to handle reducing equivalents, may impose reductive pressure and promote electron leak and superoxide generation (28,29). We therefore examined whether oxidative stress contributes to the mitochondrial defects induced by PK activation. DASA-58 increased mitochondrial superoxide levels in both WT and BRCA2-/- cells (Fig. 4G), indicating that superoxide generation is a general consequence of PK activation rather than an HRD-specific event.

However, only BRCA2-deficient cells underwent cell death upon increased superoxide levels (Fig. 4H), suggesting that the selective vulnerability lies in the capacity to tolerate this oxidative burden rather than in its generation. Consistent with this interpretation, treatment with the antioxidant glutathione, and more effectively with a membrane-permeable superoxide dismutase/catalase mimetic, suppressed DASA-58-induced cell death in BRCA2-deficient cells (Fig. 4H). Together, these findings indicate that PK activation generates superoxide in both genetic backgrounds, but BRCA2 loss converts this stress into a lethal event.

### PK activation disrupts PINK1-dependent mitophagy control selectively in BRCA2-deficient cells

Given the mitochondrial superoxide accumulation and respiratory defects induced by PK activation, we next examined PINK1, a key regulator of mitochondrial quality control and mitophagy (30). At baseline, total PINK1 abundance was markedly reduced in BRCA2-/- cells compared to BRCA2 WT cells (Fig. 5A). DASA-58 treatment did not alter PINK1 abundance but produced a mobility shift in PINK1 in BRCA2 WT cells, consistent with autophosphorylation and activation of the mitophagy program (Fig. 5A). This shift was absent in BRCA2-/- cells, indicating that PK activation engages the PINK1 axis in HR-proficient cells but fails to do so in HR-deficient cells.

**Figure 5.**
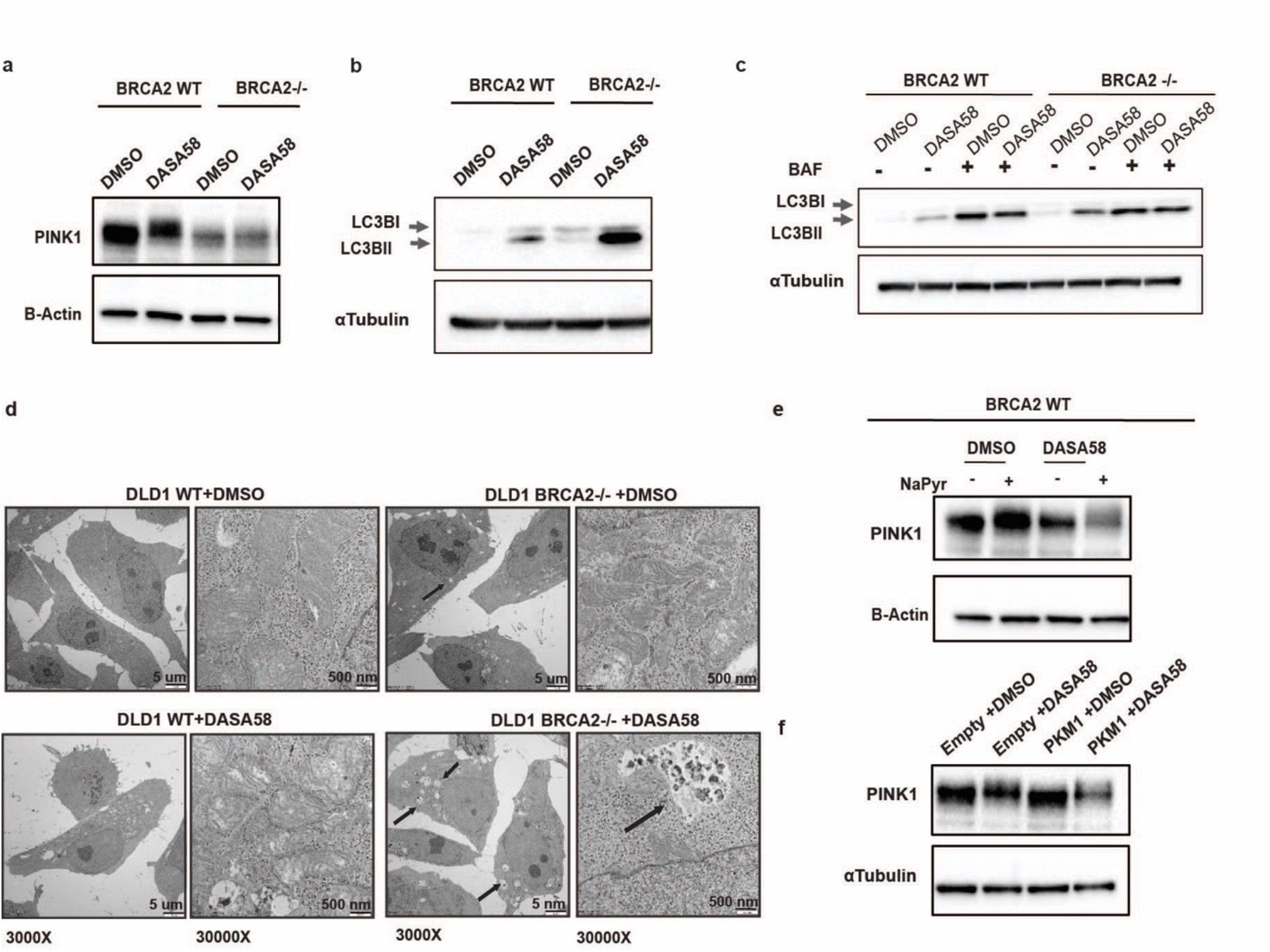
Pyruvate kinase activation disrupts PINK1-dependent mitophagy in BRCA2-deficient cells. **a** Immunoblot analysis of PINK1 in DLD1 BRCA2 WT and BRCA2-/- cells treated with DASA58 or vehicle. A representative image is shown (n=3). β-Actin was used as a loading control. **b** Immunoblot analysis of LC3B-I and LC3B-II in DLD1 WT and BRCA2-/- cells treated with or without DASA58. A representative image is shown (n=4). α-Tubulin was used as a loading control. **c** Immunoblot analysis of LC3B-I and LC3B-II in DLD1 WT and BRCA2-/- cells treated with or without DASA58, in the presence or absence of bafilomycin. A representative image is shown (n=3). α-Tubulin was used as a loading control. **d** Electron microscopy images of DLD1 WT and BRCA2-/- cells treated with or without DASA58. Arrows indicate multivesicular body (MVB)-like structures. Scale bar 5nm. Representative image are shown. **e** Immunoblot analysis of PINK1 in DLD1 BRCA2 WT cells supplemented with or without sodium pyruvate (NaPyr) and treated with or without DASA58. α-Tubulin was used as a loading control. A representative image is shown (n=3). **f** Immunoblot analysis of PINK1 in DLD1 BRCA2-/- cells transfected with empty vector or stably overexpressing PKM1, treated with or without DASA58. α-Tubulin was used as a loading control. A representative image is shown (n=4).

Consistent with these findings, DASA-58 disrupted multiple autophagy and mitophagy markers, with stronger effects in BRCA2-deficient cells (Fig. 5B-C, Supplementary Fig. S9A). LC3B-I and LC3B-II were both elevated at baseline in BRCA2-/- cells compared to WT, and DASA-58 further increased LC3B-II in both genotypes, with a more pronounced effect in BRCA2-/- cells (Fig. 5B). Bafilomycin treatment brought LC3B-II in DMSO- and DASA-58-treated samples to a similar level across both genotypes, indicating that DASA-58 raises LC3B-II by blocking its lysosomal degradation rather than by stimulating autophagosome formation (Fig. 5C). This block was accompanied by elevated p62 levels, reduced beclin-1 abundance, and decreased ULK1 phosphorylation (Supplementary Fig. S9A), each of which is consistent with impaired initiation and progression of autophagy.

Ultrastructural analysis by electron microscopy supported this conclusion. BRCA2-deficient cells treated with DASA-58 selectively accumulated multivesicular bodies (Fig. 5C, Supplementary Fig. S9B), structures whose accumulation reflects impaired endolysosomal flux and is consistent with a block at the lysosomal step of autophagic degradation.

We next asked whether these effects could be attributed specifically to the elevated PK activity state of BRCA2-/- cells, rather than to BRCA2 loss *per se*. In BRCA2 WT cells, pyruvate supplementation or DASA-58 treatment alone did not substantially alter PINK1 abundance, but the combination of pyruvate and DASA-58 strongly reduced PINK1 levels, recapitulating the baseline phenotype of BRCA2-/- cells (Fig. 5E). PKM1 overexpression in BRCA2 WT cells similarly reduced PINK1 abundance, again recapitulating the BRCA2-/- baseline (Fig. 5F). The remaining PINK1 in PKM1-overexpressing cells was still competent to undergo a DASA-58-induced shift, suggesting that the PKM1 overexpression is responsible for the PINK1 abundance reduction, but not for the defect in the activation machinery (Fig. 5F). Together, these data establish that a high PK activity state, achieved either through substrate supply combined with a PK activator or through sustained elevation of PKM1, is sufficient to reduce PINK1 abundance in HR-proficient cells, phenocopying the PINK1 deficit observed at baseline in BRCA2-deficient cells.

### The FDA-approved pyruvate kinase activator mitapivat selectively delays growth of BRCA2-deficient tumors *in vivo*

Having established that BRCA2-deficient cells are selectively vulnerable to pharmacological PK activation, we asked whether this vulnerability could be exploited therapeutically. Mitapivat is an orally available allosteric PK activator approved by the US Food and Drug Administration for the treatment of pyruvate kinase deficiency-associated hemolytic anemia (31,32), providing a clinically tractable tool to test whether PK activation can suppress HR-deficient tumors *in vivo*.

We first assessed mitapivat pharmacokinetics in mice bearing DLD1 xenografts. Following oral administration at 50, 75, and 100 mg/kg, mitapivat was detected in plasma and tumor tissue (Supplementary Fig. S10A-B), confirming tumor exposure. To verify on-target engagement, we measured PK activity in intestine, liver, blood, and tumor tissue and observed robust activation across all compartments (Fig. 6A, Supplementary Fig. 10C-E). At 50 mg/kg, mice maintained stable body weight with no change in liver or ovarian weights (Supplementary Fig. S10F-H), indicating good tolerability.

**Figure 6.**
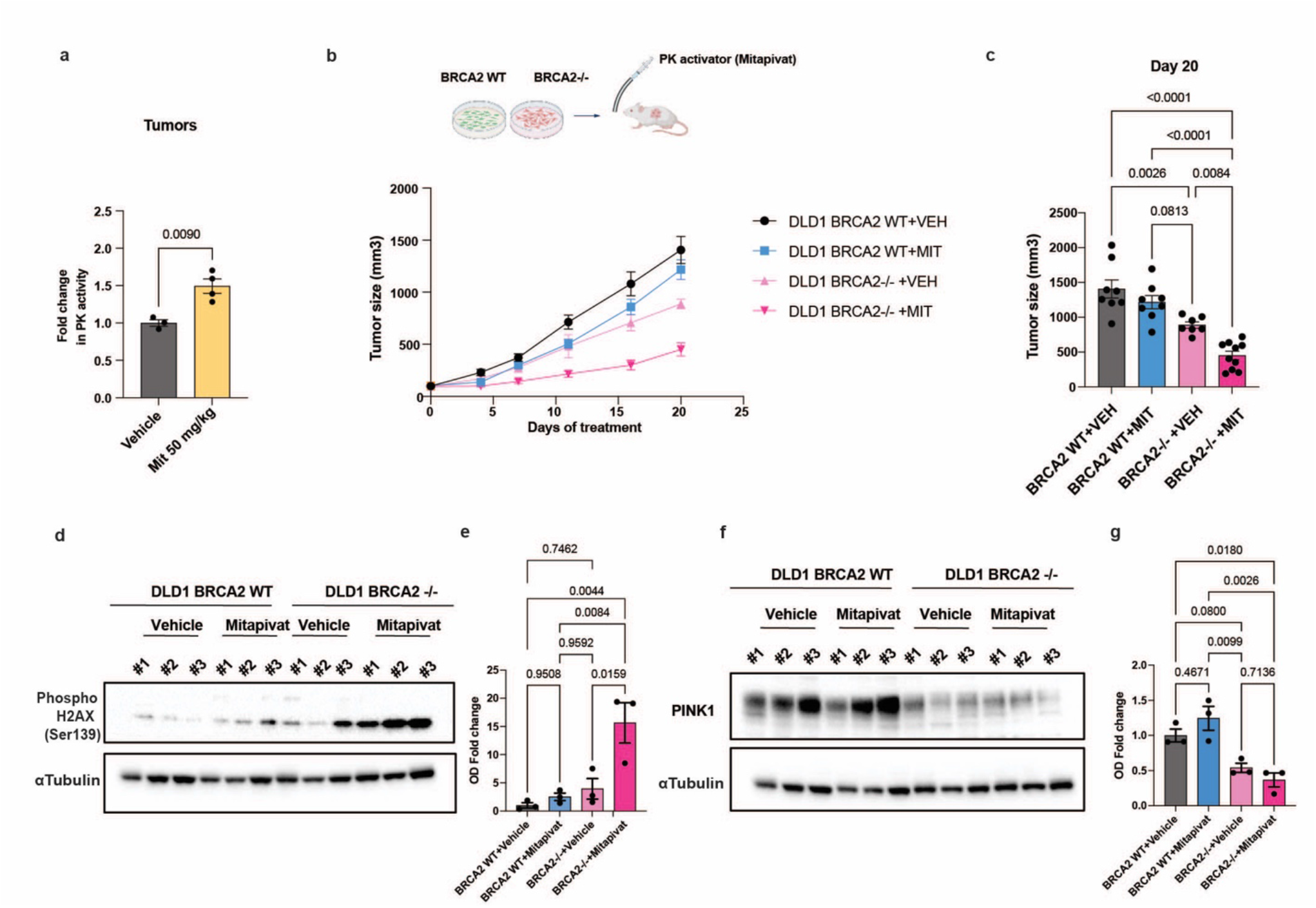
Mitapivat selectively suppresses BRCA2-deficient tumor growth in vivo. **a** PK activity assay performed on mouse tumor lysates (vehicle, n = 3; mitapivat, n = 4). **b** Schematic of the mouse experiment (created with Biorender) (schematic, top). DLD1 BRCA2 WT- and BRCA2-/--derived xenografts. Tumor volumes were measured and multiple time points are shown. **c** Tumor volumes at the final time point (day 20) (WT + vehicle, n = 8; WT + mitapivat, n = 8; BRCA2-/- + vehicle, n = 7; BRCA2-/- + mitapivat, n = 10). **d** Immunoblot analysis of phospho-H2AX (Ser139) in tumor lysates from BRCA2 WT and BRCA2-/- xenografts treated with vehicle or mitapivat; quantification is shown in **e**. α-Tubulin was used as a loading control (n=3 mice per group). **f** Immunoblot analysis of PINK1 in tumor lysates from BRCA2 WT and BRCA2-/- xenografts treated with vehicle or mitapivat; quantification is shown in **g**. α-Tubulin was used as a loading control (n=3 mice).

We next evaluated antitumor efficacy in DLD1 BRCA2 WT and BRCA2-/- xenografts (Fig. 6B). Mitapivat selectively delayed growth of BRCA2-/- tumors, while having no effect on BRCA2 WT tumors (Fig. 6C). At day 20, BRCA2-/- tumor volume was reduced by 52%, whereas BRCA2 WT tumors showed no significant difference (Fig. 6C). Thus, pharmacological PK activation with a clinically approved agent at tolerated doses is sufficient to selectively suppress HR-deficient tumor growth *in vivo*.

To assess whether the *in vivo* response reflects the same mechanism observed *in vitro*, we measured DNA damage markers and PINK1 in mitapivat-treated tumors. γH2AX was selectively elevated in BRCA2-/- tumors after treatment, consistent with the increased DNA damage seen in cultured cells (Fig. 6D-E). PINK1 abundance was reduced in BRCA2-/- tumors with and without mitapivat, mirroring the pattern seen in cell lines (Fig 6F-G).

## Discussion

Our study identifies PKM as a metabolic vulnerability in BRCA1/2-deficient cancer and reveals a therapeutic strategy to exploit it. Through integrated analysis of 10,619 TCGA tumors, isogenic BRCA1/2-deficient cell models across three lineages, and *in vivo* xenograft experiments, we find that HRD is associated with broad metabolic rewiring, with consistent alterations in glucose metabolism. Within this landscape, BRCA1/2 loss drives a selective upregulation in the constitutively active PKM1 isoform, generating an elevated PK activity state that contrasts with the PKM2-dominant phenotype of most cancers. In BRCA2-deficient cells, pushing PKM activity past a critical threshold through pharmacological overactivation reduces mitochondrial oxidative capacity and superoxide accumulation, compromises PINK1-dependent mitochondrial quality control, and culminates in selective cell death. *In vivo*, the FDA-approved PK activator, mitapivat, selectively suppresses BRCA2-deficient tumor growth while sparing HR-proficient counterparts. These findings recast HRD as a metabolic state and identify PK activation as a candidate therapeutic strategy for biomarker-selected HRD cancers.

The selective lethality of pharmacological PK activation in BRCA2-deficient cells emerges from the configuration of their pyruvate disposal machinery. BRCA2-deficient cells display reduced respiratory chain complex abundance, which limits mitochondrial oxidative capacity, and reduced LDHA protein abundance, which limits cytosolic lactate fermentation. Reduced LDHA has been previously shown to render cancer cells more sensitive to oxidative stress, with LDHA depletion in lymphoma and pancreatic cancer cells inducing ROS accumulation and cell death rescuable by N-acetylcysteine (33). While BRCA-proficient cells can buffer the extra pyruvate and reducing equivalents generated by PK activation with intact LDHA and OXPHOS capacity, BRCA2-deficient cells cannot, leading to mitochondrial overload and cell death. In this way, our findings extend prior work on context-dependent sensitivity to PKM2 activation (27) by identifying HRD as the genetic context that compresses the metabolic buffer enough to convert pharmacological PK activation from a tolerated perturbation into a lethal one.

Two complementary experiments establish that the toxic event is the entry of this additional pyruvate into mitochondria. Exogenous pyruvate sensitized BRCA2 wild-type cells to DASA-58, whereas blocking mitochondrial pyruvate import with UK5099 rescued BRCA2-deficient cells. We conclude that the pyruvate is forced into the mitochondrial compartment, where excess substrate flux through the faulty respiratory chain promotes electron leak and superoxide generation (28,29). This mechanism parallels prior demonstrations that pharmacological redirection of pyruvate into mitochondria generates mitochondrial H_2_O_2_ and triggers selective apoptosis of cancer cells (34). It is also consistent with recent work in T cells showing that genetic or pharmacological conversion of PKM2 to a constitutively active state drives mitochondrial ROS accumulation and selective cell death rescuable by mitochondrial antioxidants (35), and with the observation that PKM2 deletion in murine cardiomyocytes, leaving PKM1 as the sole isoform, produces reduced mitochondrial oxygen consumption and elevated mitochondrial superoxide (36).

Lahiguera et al. (4) similarly examined metabolic changes in BRCA-null tumors, and our results both align with and differ from theirs. The main point of agreement is LDHA: their finding of reduced *LDH* mRNA supports our observation that BRCA2-deficient cells have lower LDHA protein, consistent with impaired lactate fermentation capacity in HRD cells. The apparent difference lies in mitochondrial metabolism, as Lahiguera et al. reported increased oxygen consumption, whereas we observed reduced respiratory chain complex abundance. This discrepancy may reflect methodological differences. They measured OCR in detached cells in suspension using respirometry, while we measured adherent cells in growth media using Seahorse analysis. Their isotope tracing was also performed without glutamine or pyruvate, forcing reliance on glucose-derived TCA input, whereas our conditions included glutamine, allowing alternative anaplerotic fueling. Our data therefore suggest that BRCA2-deficient cells may preferentially route glutamine into the TCA cycle, a flexibility not captured under glutamine-free conditions. More broadly, the two studies highlight complementary vulnerabilities in HRD: Lahiguera et al. emphasize PARP/NAD+ dependence, whereas our work identifies impaired mitochondrial quality control and superoxide stress during PK activation. Together, these findings suggest that HRD cells exist within a narrow metabolic window in which either too little mitochondrial function or too much pyruvate-driven mitochondrial flux can be lethal.

PINK1 is held at low abundance on healthy, polarized mitochondria by voltage-dependent proteolysis and accumulates selectively on damaged mitochondria, where it phosphorylates ubiquitin to activate Parkin and recruit the upstream autophagy machinery (37,38). We found that total PINK1 abundance is reduced in BRCA2-deficient cells, in agreement with others who have reported that BRCA2 is directly required for Parkin-mediated clearance of damaged mitochondria (39). In WT cells, PK activation, through either pyruvate supplementation combined with DASA-58 or sustained PKM1 overexpression, was sufficient to reduce PINK1 abundance to levels comparable to those in BRCA2-/- cells. However, this reduction did not impair DASA-58-induced PINK1 activation, as measured by the characteristic PINK1 band shift. Together, these findings suggest that elevated PK activity acts upstream of PINK1 dysregulation by reducing PINK1 abundance, while loss of BRCA2 may be specifically required to disrupt the machinery responsible for PINK1 activation (39). In this way, PINK1 emerges as a proximate mediator through which the high-PK-activity state converts mitochondrial stress from a tolerable condition into a lethal signal. The mechanism by which sustained PK activity reduces PINK1 abundance, whether through accelerated turnover, suppressed expression, or altered mitochondrial localization, remains an important open question.

Mitapivat strengthens the translational relevance of our findings. Although PK activators are not established in oncology, mitapivat is already FDA-approved, orally bioavailable, and has a well-characterized safety and pharmacokinetic profile (31,32). In our models, it reached plasma and tumor tissue, activated pyruvate kinase, and selectively delayed BRCA2-deficient tumor growth at tolerated doses. Although the effect was growth delay rather than regression and tolerability testing was limited, these data provide proof-of-principle for a therapeutic window in HR-deficient tumors. Because PK activation is mechanistically distinct from PARP inhibition, it may be especially relevant in PARP inhibitor-resistant disease, and BRCA1/2 status could serve as an existing biomarker for patient selection.

This study has several limitations. Although our pan-cancer analysis suggests that HRD-associated metabolic rewiring may be broadly relevant, mechanistic work was limited to a small number of BRCA1/2-deficient models. Moreover, we did not perform genetic complementation of BRCA1/2 to reverse PKM1 upregulation; thus, strict necessity remains to be established. PKM1 upregulation upon BRCA1 loss parallels the response observed BRCA2-/- cells, supporting a shared metabolic response to HRD at the isoform level. However, the functional consequences of PK activation (cell death, DNA damage, PINK1 disruption, and *in vivo* growth inhibition) were characterized in BRCA2-deficient models, and extension of these readouts to BRCA1-deficient models represents an immediate next step. A key caveat is that our functional and *in vivo* therapeutic experiments were performed exclusively in BRCA2-deficient models. Given that BRCA1 and BRCA2 act at distinct steps of HR, we cannot assume equivalent PK activator sensitivity in BRCA1-deficient cells despite the shared PKM1 upregulation phenotype. Therefore, it remains unclear whether PKM1 upregulation and PK activator sensitivity extend to other HR-deficient genotypes such as those lacking RAD51, PALB2, or Fanconi anemia pathway genes. The mechanism of PKM1 induction after BRCA1/2 loss also remains unresolved. We did not detect transcriptomic changes in canonical regulators of PKM splicing suggesting that this process may be mediated by post-translational modifications and/or additional, as-yet-unidentified factors. In addition, key elements of the proposed mechanism, including shuttle activity, compartmental NAD+/NADH balance, and the basis of PINK1 loss, were not directly measured. Finally, our *in vivo* studies were limited and do not yet address how lineage, microenvironment, co-occurring mutations, or chronic treatment influence response.

Despite these caveats, our findings have broader implications for cancer metabolism. They show that genotype-specific metabolic states can create actionable vulnerabilities, and that isoform-level rewiring may be more informative than total pathway activity. PKM1 emerges as a candidate biomarker for sensitivity to PK activation, pending validation in larger cohorts. More broadly, our work identifies PK as a metabolic node coupling HRD to mitochondrial fragility and identifies a therapeutic strategy that exploits this coupling. The convergence of an actionable metabolic vulnerability, a clinically approved drug, and an established patient-stratifying biomarker positions PK activation as a strategy worthy of clinical evaluation in HRD cancers and, more broadly, motivates a view of metabolic dependencies as emergent properties of the DNA repair landscape.

## METHODS

### Differential gene expression analyses across cancer types

Gene-level RNA-sequencing unstranded count matrices for primary tumours were obtained from The Cancer Genome Atlas (TCGA) via TCGAbiolinks (v2.34.0) (20) across all available cancer types. Homologous recombination deficiency (HRD) scores were sourced from the pan-cancer analysis of Shi et al. (21), in which HRD was quantified using a composite score derived from loss of heterozygosity (LOH), large-scale state transitions (LST) and telomeric allelic imbalance (NtAI). Analyses were restricted to cancer types for which both HRD scores and RNA-seq count data were available. Count matrices were library-size normalized and filtered using filterByExpr with default parameters, followed by trimmed mean of M-values normalization using edgeR (v4.4.0). Differentially expressed genes were identified using the limma-voom pipeline (v3.62.1), with HRD score included as a continuous variable in the design matrix. Analyses were performed independently within each cancer type.

### Differential Protein Expression Analyses Across Cancer Types

Somatic mutation data were obtained from the Clinical Proteomic Tumor Analysis Consortium (CPTAC) (22) cohorts TCGAbiolinks (v2.34.0). The proteomics data in log2 ratios was downloaded directly using the proteomics data commons graphql endpoint (https://pdc.cancer.gov/graphql). Tumor types included clear cell renal cell carcinoma (ccRCC; PDC000127), colon adenocarcinoma (COAD; PDC000116), head and neck squamous cell carcinoma (HNSCC; PDC000221), lung adenocarcinoma (LUAD; PDC000153), lung squamous cell carcinoma (LSCC; PDC000234), pancreatic ductal adenocarcinoma (PDAC; PDC000270), and uterine corpus endometrial carcinoma (UCEC; PDC000125, PDC000439). Protein abundance measurements were derived from CPTAC normalized protein expression matrices reported as log2 abundance ratios. For tumor types with multiple CPTAC proteomic batches, abundance matrices were merged into a single matrix retaining only protein abundances shared across batches. Somatic mutation data were obtained from CPTAC whole-exome sequencing mutation annotation files (MAFs). For colon adenocarcinoma, mutation calls and sequencing metadata were obtained from the corresponding cBioPortal CPTAC release. Samples harboring at least one qualifying loss-of-function (LoF) mutation in BRCA1, BRCA2, PALB2, RAD51 (HRD genes) were classified as homologous recombination deficient (HRD). Tumors containing only non-LoF mutations in HRD genes were excluded from analysis. Differentially expressed proteins were identified using the limma (v3.62.1), with HRD status included as a binary variable in the design matrix. Analyses were performed independently within each cancer type.

### Pathway analyses

Gene lists from limma differential expression analysis were pre-ranked by log fold change (logFC; RNA-seq) or t-statistic (proteomics) and used as input for gene set enrichment analysis (GSEA). Enrichment of Hallmark and C2 KEGG gene sets (MSigDB; retrieved using msigdbr v10.0.1) was tested using fgsea (v1.32.2) with parameters minSize = 15, maxSize = 500 and nPermSimple = 10,000. Multiple testing correction was applied using the Benjamini–Hochberg method, and pathways with an adjusted *P* < 0.05 were considered statistically significant. All analyses were performed in R (v4.4.3). Data manipulation and visualization were performed using tidyverse (v2.0.0), cowplot (v1.1.3), and ggrepel (v0.9.6).

### Cell lines and culture conditions

DLD1 BRCA2 WT and DLD1 BRCA2−/− cells were purchased from Horizon Discovery (HD 105-007). ID8 Trp53−/− Brca2 WT and Brca2−/− cells were kindly provided by the McNeish laboratory. K8082 pancreatic cells were kindly provided by the Kenneth Olive laboratory. All cell lines were maintained in Dulbecco’s Modified Eagle Medium (DMEM; Thermo Fisher Scientific, Cat:#11965092) supplemented with 10% fetal bovine serum (R&D Systems, Cat:#11150). Cells were routinely tested for mycoplasma contamination and maintained at 37 °C in a humidified incubator with 5% CO_2_.

### Drugs and reagent preparation

DASA-58 was purchased from MedChemExpress (Cat:#HY-19330). UK5099 was purchased from Selleck Chemicals (Cat:#PF-1005023). Reduced glutathione (GSH) was purchased from MilliporeSigma (Cat:#G6013) and prepared fresh for each experiment. MnTMPyP, a superoxide dismutase mimetic, was purchased from MilliporeSigma (Cat:#475872) and dissolved in sterile water. Stock solutions were stored at −20 °C. Bafilomycin was purchased from Santa Cruz Biotechnology (Cat:#sc-201550). For *in vivo* experiment Mitapivat was purchased from STARK Chemicals (Cat:#1260075-17-9) and dissolved in 5% DMSO, 40% PEG400, 5% Tween-80 and 50% sterile water. Mitapivat was freshly prepared for each treatment.

### Viability assays

For cell death experiments, 90,000 DLD1 BRCA2 WT, 117,000 DLD1 BRCA2−/−, and 40,000 ID8 Trp53−/− Brca2 WT or Brca2−/− cells were seeded in 6-well plates. DLD1 BRCA2 WT and BRCA2−/− cells were plated at different densities to account for differences in growth rate such that both genotypes reached similar confluence at the time of treatment, 24 h after seeding. At 24 h after plating, medium was replaced with fresh medium containing vehicle (DMSO) or DASA-58 (50 μM). Cells were treated for 92 h (DLD1) or 48 h (ID8), with treatment duration adjusted according to the growth characteristics of each cell line. For antioxidant rescue experiments, GSH (5 mM) and MnTMPyP (500 μM) were added together with DASA-58. Cells number was determined by using Trypan blue/Erythrosine B exclusion assay using a Neubauer hemocytometer.

### RNA interference with siRNA oligonucleotides

DLD1, ID8 Trp53−/− and K8082 cells were transfected using Lipofectamine RNAiMAX (Thermo Fisher Scientific, Cat:#13778030) according to the manufacturer’s instructions. Cells were seeded the day before transfection in 6-well plates (DLD1, 200,000 cells per well; ID8 Trp53−/− and K8082, 85,000 cells per well) and transfected at 40–50% confluence with the indicated siRNA oligonucleotides (25 nM for human BRCA2 and 50 nM for mouse Brca2) using Lipofectamine RNAiMAX (Thermo Fisher Cat:# 13778100). Cells underwent two rounds of transfection, the first at seeding and the second 24 h later. The following oligonucleotides were used: ON-TARGETplus Human BRCA2 siRNA (Cat:#L-003462-00-0005), ON-TARGETplus Mouse Brca2 siRNA (Cat:#L-042993-00-0005), ON-TARGETplus Human BRCA1 siRNA (Cat:#L-003461-00-0005) (Horizon Discovery) and ON-TARGETplus non-targeting control pool (Cat# L2-020459-01-0005).

### PKM1 overexpression

pLHCX-FLAG-mPKM1 was purchased from Addgene (plasmid#44240). pLHCX-FLAG empty backbone was generated by BamHI/ClaI digestion to remove the mPKM1 insert, followed by Klenow-mediated blunting of digested ends, and T4 DNA ligation. Retroviral particles were generated in HEK-293 cells transfected with pLHCX-FLAG-mPKM1 or pLHCX-FLAG empty (10µg), pCL-Eco (3 µg), and VSVG (1 µg) plasmids. Virus-containing supernatant from transfected cells was collected at 48h and 72h after transfection, pooled, centrifuged, and filtered through a 0.45 µm filter. Viral supernatants were stored at -80° C and used to infect target cells (200,000 cells for DLD1 BRCA2 WT, 85,000 cells for ID8 Trp53-/-) in a 6-well plate in the presence of polybrene (4 μg/ml). Successfully infected cells were selected using hygromycin (0.25 mg/ml for DLD1 and 1mg/ml for ID8).

### PKM knockdown

pLKO-shPKM1/2 (pLKO-shPKM2_4) was a gift from D. Anastasiou (Addgene plasmid #42516), and scramble shRNA was a gift from D. Sabatini (Addgene plasmid #1864). Lentiviruses were produced in HEK293T cells by co-transfection of plasmids expressing Gag/Pol, Rev, and VSV-G together with the respective pLKO vector (10 µg). Virus-containing supernatant was collected 48 and 72 h after transfection, centrifuged to remove cell debris, filtered through a 0.45 µm filter, aliquoted, and stored at −80°C until use. For knockdown experiments, DLD1 BRCA2 wild-type and DLD1 BRCA2−/− cells were seeded at 300,000 cells per well in 6-well plates and infected with viral supernatant in the presence of 8 µg/mL polybrene. Cells were selected with 2 µg/mL puromycin for 2 weeks to generate stable pooled shRNA-expressing populations. PKM1/2 knockdown was confirmed by immunoblotting. For cell confluence experiments, puromycin-selected DLD1 BRCA2 wild-type and DLD1 BRCA2−/− cells expressing either shPKM1/2 or scramble shRNA were seeded in 96-well plates at 3,000 cells per well, with 8 replicate wells per condition. Cells were maintained in medium containing 2 µg/mL puromycin throughout the experiment. Unused perimeter wells were filled with PBS or medium to minimize edge effects. Cell confluence was measured using an IncuCyte SX5 Live-Cell Analysis System (Sartorius) according to the manufacturer’s instructions. Phase-contrast images were acquired, and percent confluence was quantified using IncuCyte analysis software. For viability assay, cells were seeded at 200,000 cells per well in 6-well plates. Cells number was determined by using Erythrosine B exclusion assay using a Neubauer hemocytometer. Data are presented as mean confluence ± SEM across replicate wells.

### RNA sequencing and analysis

DLD1 BRCA2 WT and BRCA2 Cells were cultured in DMEM and treated with DASA-58 (50 μM) or vehicle (DMSO) for 48 h. Total RNA was extracted using the RNeasy kit (Qiagen). For library preparation and sequencing, 2.5 μg purified RNA was submitted to GENEWIZ NGS Services from Azenta Life Sciences (South Plainfield, NJ, USA). Libraries were prepared using poly(A) selection and sequenced on an Illumina platform (Illumina NovaSeq X Plus instrument) using a 2 × 150 bp configuration. Raw sequencing reads (FASTQ files) were aligned to the human reference genome (Homo sapiens GRCh38.110) using HISAT2. Read counts were generated using featureCounts (Subread v2.0.1) in RStudio (2023.06.1). Gene set enrichment analysis (GSEA), including nominal P values, adjusted P values and normalized enrichment scores (NES), was performed using the R package fgsea (v1.32.4). Graphs were generated in RStudio (2023-10-31) with R (v4.3.2) using ggplot2 (v4.0.3).

### RNA extraction, RT-PCR and qRT-PCR

Total RNA was isolated using the RNeasy kit (Qiagen) according to the manufacturer’s instructions. Briefly, 1 μg purified RNA was reverse-transcribed using SuperScript VILO Master Mix (Thermo Fisher Scientific, Cat:#11756050). Resulting cDNA (1:20, v/v) was analysed by quantitative real-time PCR using PowerUp SYBR Green Master Mix (Thermo Fisher Scientific, Cat:#A25742) on a QuantStudio 3 Real-Time PCR System. Primers targeting human and mouse BRCA2/Brca2, PKM1 and PKM2 were used. Transcript levels were normalized to GAPDH. Primer sequences were as follows:

**Mouse Brca2**

Forward: CGAAGTCAAACTCTACCACTGGA

Reverse: CCACAGGGTCCACTTTGGTC

**Human BRCA2**

Forward: ACTCTGCCGCTGTACCAATC

Reverse: AGTTTTCACTGTGCGAAGACTTT

**Mouse PKM1**

Forward: GTCTGGAGAAACAGCCAAGG

Reverse: TCTTCAAACAGCAGACGGTG

**Human PKM1**

Forward: CAGCCAAAGGGGACTATCCT

Reverse: GAGGCTCGCACAAGTTCTTC

**Mouse PKM2**

Forward: GTCTGGAGAAACAGCCAAGG

Reverse: CGGAGTTCCTCGAATAGCTG

**Human PKM2**

Forward: GTGGGGTCGCTGGTAATG

Reverse: TTCCTCAAATAATTGCAAGTGG

**Human and mouse GAPDH**

Forward: TCAAGAAGGTGGTGAAGCAGG

Reverse: ACCAGGAAATGAGCTTGACAAA

Relative gene expression was calculated using the 2^−ΔCt^method.

### Mitochondrial DNA copy number

Total DNA was isolated from DLD1 cells using the DNeasy Blood and Tissue Kit (Qiagen) and treated with RNase A according to the manufacturer’s protocol. Mitochondrial DNA content (ND1) relative to nuclear DNA (GAPDH) was quantified by real-time PCR using a QuantStudio 3 Real-Time PCR System. Primer sequences were as follows:

**ND1**

Forward: TCTCCACCCTTATCACAACA

Reverse: GACTAGTTCGGACTCCCCTT

**GAPDH**

Forward: CTCCTGTTCGACAGTCAGC

Reverse: TTCAGGCCGTCCCTAGC

Mitochondrial DNA copy number was calculated using the 2 × 2^−ΔCt^ method, as previously described ^43^.

### Protein extraction and western blot analysis

Cells were cultured in DMEM and treated with DASA-58 (50 μM) or vehicle (DMSO) for 48 h (DLD1) or 24 h (ID8). For sodium pyruvate rescue experiments, cells were co-treated with 10 mM sodium pyruvate (Thermo Fisher Scientific, Cat:#11360070). For autophagy-related experiments, cells were treated with bafilomycin (1 μM) for 6 h before lysis. At harvest, cells were washed twice with ice-cold PBS and lysed in RIPA buffer supplemented with 0.01% SDS (0.5 M Tris-HCl, pH 7.4, 1.5 M NaCl, 2.5% deoxycholic acid, 10% NP-40, 10 mM EDTA; MilliporeSigma, Cat:#20188), protease inhibitor cocktail (MilliporeSigma, Cat:#P8340) and phosphatase inhibitors (PhosSTOP, Roche, Cat:#4906845001). Tumour tissues were collected immediately after euthanasia, snap-frozen in liquid nitrogen and stored at −80 °C until use. For mouse tumour lysates, tumours were homogenized using a TissueLyser III (Qiagen) in RIPA buffer supplemented with protease and phosphatase inhibitors, followed by centrifugation at 21,000g for 30 min. Supernatants were collected and centrifuged again at 21,000g for 20 min, and the final supernatants were used for analysis. Protein concentration was determined using a BCA assay (Thermo Fisher Scientific, Cat:#23225). Proteins were resolved by SDS–PAGE and analyzed by immunoblotting using the following antibodies: BRCA2 (1:1,000, Cell Signaling Technology, Cat:#10741), BRCA1 (1:1,000, Cell Signaling Technology, Cat:#9010S), PKM1/2 (1:1,000, Cell Signaling Technology, Cat:#3190), HK1 (1:1,000, Cell Signaling Technology, Cat:#2024S), LDHA (1:1,000, Cell Signaling Technology, Cat:#2012S), PFK (1:1,000, Cell Signaling Technology, Cat:#8164), PDH (1:1,000, Cell Signaling Technology, Cat:#3205S), phospho-PDH Ser293 (1:1,000, Cell Signaling Technology, Cat:#37115), PKM1 (1:1,000, Cell Signaling Technology, Cat:#7067), PKM2 (1:1,000, Cell Signaling Technology, Cat:#4053), phospho-H2AX Ser139 (γH2AX; 1:5,000, Millipore, Cat:#05-636), PINK1 (1:1,000, Cell Signaling Technology, Cat:#6946), α-tubulin (1:10,000, Cell

Signaling Technology, Cat:#3873), β-actin (1:5,000, Cell Signaling Technology, Cat:#4970), LC3B I/II (1:1,000, Cell Signaling Technology, Cat:#3868), p62 (1:1,000, Cell Signaling Technology, Cat:#8025), Beclin1 (1:1,000, Cell Signaling Technology, Cat:#3495), ULK1 (1:1,000, Cell Signaling Technology, Cat:#8054), phospho-ULK1 Ser555 (1:1,000, Cell Signaling Technology, Cat:#5869), OxPhos Rodent WB Antibody Cocktail (1:1,000, Abcam, Cat:#ab110413), TIM23 (1:1,000, Cell Signaling Technology, Cat:#34822), TOM20 (1:1,000, Cell Signaling Technology, Cat:#42406) and Vinculin (1:1,000, Cell Signaling Technology, Cat:#4650). Immunoreactive bands were detected using a ChemiDoc imaging system (Bio-Rad) with chemiluminescent substrates (Pierce ECL Western Blotting Substrate and SuperSignal West DURA, Thermo Fisher Scientific, Cat:#32109 and Cat:#34075 respectively). Band intensities were quantified using Image Lab software.

### Seahorse analysis

Oxygen consumption rate (OCR) was measured using a Seahorse XFe24 Analyzer (Agilent). DLD1 cells were seeded in Seahorse 24-well assay plates the day before analysis. DLD1 BRCA2 WT and BRCA2−/− cells were plated at 40,000 and 50,000 cells per well, respectively. DLD1 BRCA2 WT and BRCA2−/− cells were plated at different densities to account for differences in growth rate such that both genotypes reached similar confluence at the time of Seahorse analysis, 24 h after seeding.

### 13C Tracing experiments and metabolomics

For isotope tracing experiments, cells were cultured in DMEM (Thermo Fisher Scientific, Cat:#11965092) supplemented with 10% FBS (R&D Systems, Cat:#11150) and treated with DASA-58 (50 μM) or vehicle (DMSO) for 48 h. At 6 h, 2 h or 30 min before the end of treatment, cells were washed twice with PBS and medium was replaced with glucose- and glutamine-free, phenol red-free DMEM (Thermo Fisher Scientific, Cat:#A1443001) supplemented with 25 mM ^12^C-glucose (Thermo Fisher Scientific, Cat:#A2494001) or uniformly labelled ^13^C-glucose (Cambridge Isotope Laboratories, Cat:#CLM-1396), 10% dialyzed FBS (R&D Systems, Cat:#11150) and 4 mM glutamine (Thermo Fisher Scientific, Cat:#25030081). Cells were lysed in 80% methanol for metabolite extraction after 6 h (citrate labelling), 2 h (malate labelling) or 30 min (lactate labelling). Polar metabolites were analyzed by z-HILIC mass spectrometry. Isotopic labelling incorporation was calculated using Mass Hunter software

(Agilent). For steady-state metabolomics, cells were treated with DASA-58 (50 μM) or vehicle (DMSO) for 1 h before lysis. Cells were lysed in 80% methanol containing 1% formic acid, and samples were submitted to NYU Langone for metabolite extraction. Polar metabolites were analysed using p-HILIC chromatography. Metabolomics data were analysed using Xcalibur software and normalized to total metabolite abundance.

### Electron microscopy

DLD1 cells were treated with DASA-58 (50 μM) or vehicle (DMSO) for 48 h and then fixed in EM “yellow” fix (2.5% glutaraldehyde, 4% paraformaldehyde, 0.02% picric acid in 0.1 M sodium cacodylate buffer, pH 7.3) overnight at 4 °C. Samples were processed and embedded by the Weill Cornell Medicine Electron Microscopy Core Facility. Images were acquired using a JEOL 1400 transmission electron microscope. The number of MVBs per cell was manually counted in 9–12 cells for each experimental group.

### Immunohistochemistry

Immunohistochemistry was performed on formalin-fixed, paraffin-embedded tissues. Slides were deparaffinized in Histo-Clear and rehydrated through a graded ethanol series to water. Antigen retrieval was performed in sodium citrate buffer (0.01 M citrate, 0.05% Tween-20, pH 6.0) using a pressure cooker for 10 min, followed by one wash in PBS containing 0.05% Tween-20 and quenching of endogenous peroxidase activity with 3% H_2_O_2_ for 10 min. Sections were blocked in 5% goat serum in PBS containing 0.05% Tween-20 for 1 h and incubated with primary antibodies overnight at 4 °C. PKM1 (Cell Signaling Technology, Cat#7067) was used at 1:200 dilution and PKM2 (Cell Signaling Technology, Cat#4053) at 1:300 dilution. Detection was performed using the DAB detection kit (Vector Laboratories, Cat#SK-4100) according to the manufacturer’s instructions, followed by hematoxylin counterstaining. Scanned H&E images were downloaded from HistoWiz as ScanScope Virtual Slide (SVS) files. Signal quantification was performed using QuPath.

### Pyruvate Kinase activity assay

To measure pyruvate kinase (PK) activity, cells were lysed in PK lysis buffer (50 mM Tris-HCl, pH 7.5, 1 mM EDTA, 150 mM NaCl, 1% IGEPAL CA-630, 50 mM HEPES) supplemented with protease inhibitors (MilliporeSigma, Cat#P8340). For mouse tumor and intestine and liver lysates, tissues were homogenized using a TissueLyser III (Qiagen) in PK lysis buffer supplemented with protease inhibitors, followed by centrifugation at 21,000g for 30 min. Supernatants were collected, centrifuged again at 21,000g for 20 min, and the final supernatants were retained. Samples were treated with HemoGlobin Depletion reagent (HemogloBind; Biotech Support Group, Cat#H0145-05) to remove hemoglobin contamination. Blood was collected by cardiac puncture. Protein concentration was determined using a BCA assay (Thermo Fisher Scientific, Cat#23225), and 2 μg total protein was used per assay. Liver lysates were normalized by weight, whereas blood was normalized by RBC counting.PK activity was measured in cell and tissue lysates using a previously described lactate dehydrogenase (LDH)-coupled assay in which phosphoenolpyruvate (PEP, 0.5 mM) is converted by PK to pyruvate, followed by rapid conversion of pyruvate to lactate by LDH (40). LDH-dependent NADH consumption was monitored kinetically by absorbance at 340 nm using a microplate spectrophotometer (BMG Labtech or Varioskan LUX, Thermo Scientific). To distinguish PKM1 from PKM2 activity, lysates were pre-incubated with 5 mM alanine for 10 min at 37 °C, as previously described (41). Activity was measured in cell or tissue lysates and the percentage of maximal activity in BRCA2 WT vs BRCA2-/- with or without PK activators was calculated. PEP concentration in the final reaction well was 0.5mM for all the experiments.

### Mitochondrial superoxide measurement

Mitochondrial superoxide was measured using MitoSOX Red (Invitrogen, Cat#M36008). MitoSOX Red, dissolved in DMSO, was added to cell suspensions and incubated at 37 °C for 20 min at a final concentration of (1μM). Cells were then washed three times with PBS and counterstained with DAPI (0.5 μg/ml). Samples were analyzed on a Sony ID7000 spectral cell analyzer using appropriate excitation and emission settings. Gating strategy is shown in Supplementary Fig. 1).

### Mouse experiments

Female nude mice (6–8 weeks old) were obtained from The Jackson Laboratory. Animals were housed under specific pathogen-free conditions at 22 ± 2 °C, 55 ± 10% relative humidity and a 12 h light–dark cycle. All animal studies were approved by the Institutional Animal Care and Use Committee (IACUC) of Weill Cornell Medical College and conducted in accordance with institutional guidelines. For xenograft studies, 5 × 10^6^ DLD1 BRCA2 WT or DLD1 BRCA2−/− cells resuspended in 100 μl PBS were injected subcutaneously in the right flank of female nude mice. Once tumors reached 100 mm^3^ (approximately 7 d after inoculation), mice were randomly assigned to treatment groups. Body weight was recorded weekly and tumor volume was measured every 2–3 d using digital calipers and calculated as (length × width^2^) × 0.5, where length and width are expressed in millimeters. Mice were treated with mitapivat (50 mg/kg) or vehicle by oral gavage (o.g.) twice daily for 20 days. Mice showing signs of pain or distress, ulcerated tumors or body weight loss greater than 20% were euthanized. At study endpoint, mice were euthanized by CO_2_ inhalation, and tissues were collected for downstream analyses, including tumors, intestine, liver, ovaries and oviducts. Tumors were divided into four equal pieces and snap frozen. Small intestines were removed, flushed with cold PBS, cut into pieces and snap frozen. Liver and ovaries/oviducts were weighed for toxicity assessment. Blood was collected by cardiac puncture.

### Mitapivat bioavailability study

Mitapivat concentrations were measured in mouse plasma and tumor samples. Tumor-bearing mice were treated with mitapivat by oral gavage at different doses (vehicle, 50, 75 or 100 mg/kg) twice daily for 5 days. Two hours after the final dose, plasma was collected from blood obtained by cardiac puncture and subcutaneous tumors were harvested and snap frozen. Blood samples were collected in EDTA-containing tubes, gently mixed to ensure anticoagulation, and centrifuged at 3,000g for 10 min at 4 °C. Samples were analyzed using a targeted LC–MS assay for mitapivat after normalization of metabolite extraction to sample input (10 mg/ml for tumor samples and 50 μl/ml for plasma samples). A seven-point standard curve (10 μM to 10 nM) of neat mitapivat was randomized and run in duplicate with each batch. Metabolite peak intensities were extracted according to a library of m/z values and retention times established using authentic standards. Intensities were extracted using an in-house script with a 10ppm tolerance for theoretical m/z and a maximum retention time window of 30 s.

### Statistical analysis

GraphPad Prism (v10) was used for statistical analyses and graphical representation. Comparisons between two groups were performed using two-sided unpaired Student’s *t*-tests. Comparisons among multiple groups were performed using one-way ANOVA followed by Tukey’s multiple-comparisons test or two-way ANOVA. *P* ≤ 0.05 was considered statistically significant. Data are presented as mean ± S.E.M. from at least three independent experiments.

## Data availability

All data supporting the findings of this study are available within the article and its Supplementary material. All other data supporting the findings of this study are available from the corresponding author on reasonable request. Materials that are subject to existing intellectual property obligations will be made available upon execution of a material transfer agreement. Source data are provided with this paper.

## Acknowledgements

This work was supported in part by MSK SPORE (found #5290123101) (to M.D.G.), the AIRC-Italian Association for Cancer Research-Francesca Barbieri Fellowship (to M.D.T.) and AICF-Italian American Cancer Foundation (to M.D.T.). We acknowledge NYU Langone Health’s Metabolomics Laboratory (RRID: SCR_017935) for its help in acquiring and analyzing the data presented. The authors would like to thank the Electron Microscope Core Laboratory in WCM. We thank K. Olive and I. McNeish for sharing cell lines with Cantley Laboratory. We also thank A. Brambati and A. Marzio for advice, discussion and for sharing reagents.

## Author contributions

M.D.G. and M.D.T. conceived the study. M.D.G, L.C.C. and M.D.T. contributed to study conceptualization. M.D.T. performed in vitro and in vivo experiments. M.A. and S.L. contributed to in vivo experiment. M.L., C.E, I.N., J.S., J.K., E.D. and D.P. contributed to in vitro experiments. N.R, E.T and E.R. contributed to TCGA and CPTAC-based data. A.S. and K.R. contributed to metabolomic-tracing data. All authors evaluated the results and edited the manuscripts. M.D.G. and M.D.T. wrote the manuscript with inputs from all the authors.

## Notes

### Competing Interest Statement

M.D.G. holds equity in Faeth Therapeutics and Skye Biosciences; reports consulting or advisory roles with Almac Discovery, Faeth Therapeutics, Genentech Inc., Scorpion Therapeutics, Skye Biosciences, and Third Arc Bio, Inc.; patents, royalties, and other intellectual property with Weill Cornell Medicine and Faeth Therapeutics. L.C.C. is a cofounder and member of the Scientific Advisory Board (SAB) and holds equity in Faeth Therapeutics, which focuses on dietary intervention during cancer therapy, Volastra Therapeutics, and Larkspur Therapeutics. He is also a cofounder, former member of the SAB and holds equity in Agios Pharmaceuticals and Petra Pharmaceuticals (now owned by Loxo@Lilly).

