## Supplementary Information for "BRCA2 loss drives a PKM1-dominant glycolytic state that is selectively lethal to pyruvate kinase activation"

1 **Supplementary Information**

a

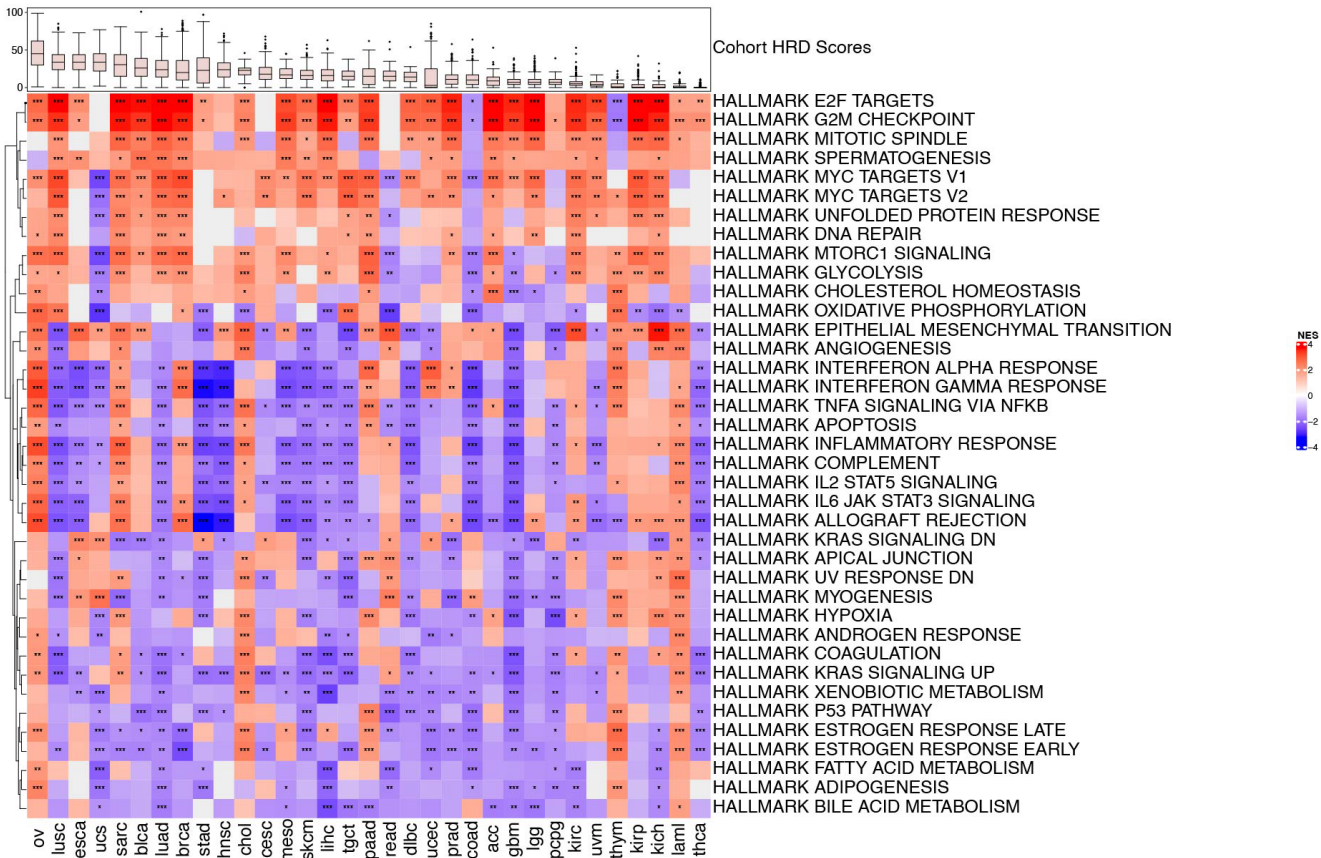

3 **Supplementary Fig.1. Differential gene expression analysis in human pan-cancer tissues.**

4 **a** Heatmap showing normalized enrichment scores (NES) for significantly enriched Hallmark gene
5 sets performed across 10,619 TCGA cancer types stratified by HRD score. Cell color denotes the
6 NES, with red indicating positive enrichment and blue indicating negative enrichment. Positively
7 enriched pathways (red) are progressively upregulated with higher HRD score, whereas
8 negatively enriched pathways (blue) are progressively downregulated. Color intensity reflects the
9 relative magnitude of regulation. All Hallmark gene sets are displayed.

a

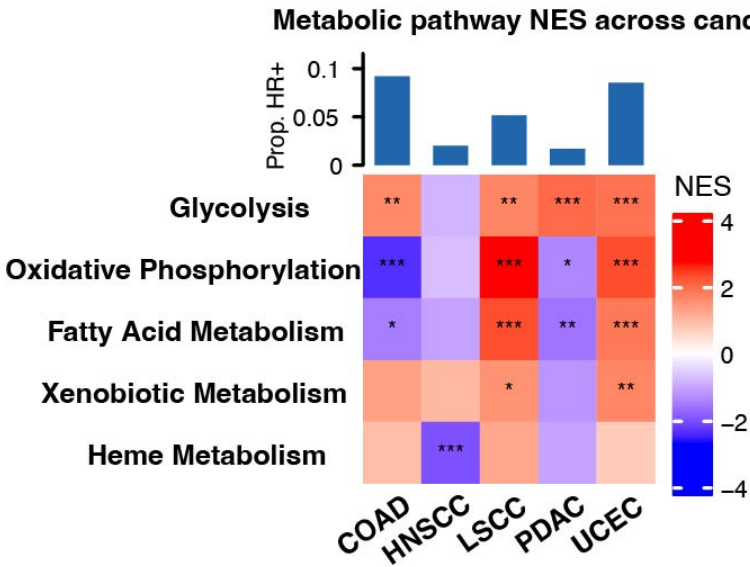

**Supplementary Fig.2. Differential protein expression analysis in human cancer.** a Heatmap showing normalized enrichment scores (NES) for significantly enriched Hallmark pathways on human cancer stratified according to the presence of at least one qualifying loss-of-function (LoF) mutation in BRCA1, BRCA2, PALB2, RAD51 (HRD genes). Tumors with enough sample size to perform differential expression analysis were analyzed and shown. Cell color denotes the NES, with red indicating positive enrichment and blue indicating negative enrichment. Positively enriched pathways (red) are upregulated HRD tumors, whereas negatively enriched pathways (blue) are downregulated. Color intensity reflects the relative magnitude of regulation. All Hallmark gene sets are displayed.

a

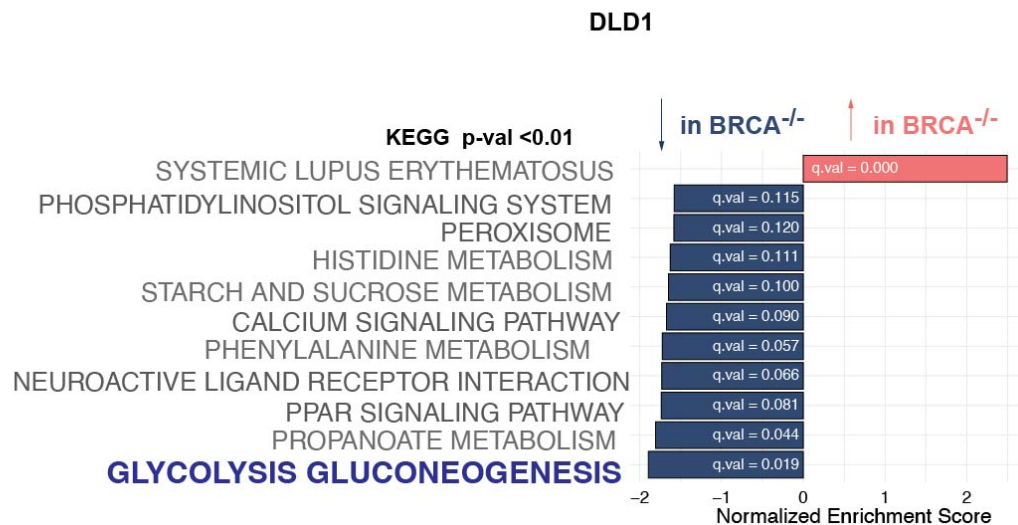

**Supplementary Fig.3. Differential gene expression analysis in DLD1 tumor cells. a** Gene set enrichment analysis (GSEA) comparing DLD1 BRCA2 WT vs BRCA2<sup>-/-</sup> cells. Upregulated pathways (red) and downregulated pathways (blue) ranked by normalized enrichment score are shown.

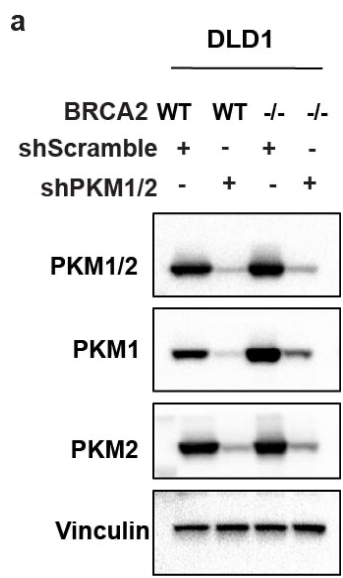

**Supplementary Fig.4. Stable PKM1/2 knockdown in DLD1 BRCA2 WT and BRCA2<sup>-/-</sup> tumor cells. a** Immunoblot analysis of PKM1/2, PKM1, PKM2 in DLD1 BRCA2 WT and BRCA2<sup>-/-</sup>

infected with sh*Scramble* or sh*PKM1/2*. Vinculin as loading control. A representative image of two-independent experiment is shown.

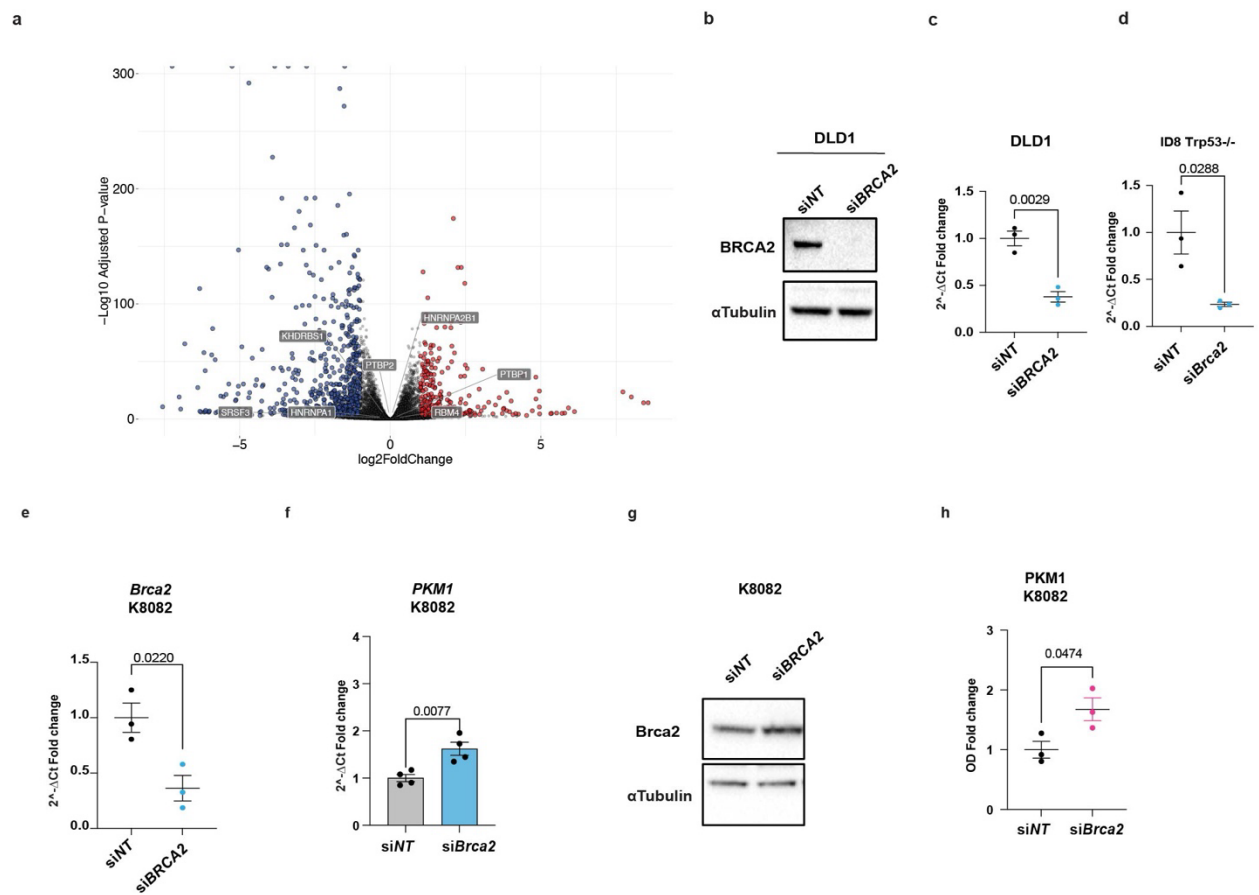

**Supplementary Fig. 5. Transient or stable loss of BRCA2 results in PKM1 upregulation without changes in the transcriptional expression of known PKM splicing regulators.**

**a** Volcano plot showing differential gene expression analysis from RNA-seq data. Each point represents a gene, plotted according to log<sub>2</sub> fold change (x-axis) and -log<sub>10</sub> adjusted *p*-value (y-axis). Significantly downregulated genes are shown in blue, significantly upregulated genes are shown in red, and genes that did not change significantly are shown in grey. KHDRSB1, PTBP1, PTBP2, RBM4, SRSF3, HNRNPA1, HNRNAA2B1 are highlighted. **b** Immunoblot analysis of BRCA2 in DLD1 BRCA2 WT cells transiently transfected with non-targeting siRNA (siNT) or siRNA targeting BRCA2 (siBRCA2). A representative image is shown. BRCA2 mRNA levels quantified by qPCR in DLD1 cells (**c**), ID8 Trp53<sup>-/-</sup> cells (**d**) and K8082 cells (**e**). **f** PKM1 mRNA levels quantified by qPCR in K8082 cells transfected with non-targeting siRNA (siNT) or siRNA

targeting Brca2 (siBrca2). **g** PKM1 protein levels detected by immunoblot. Quantification of immunoblots shown in **h**.

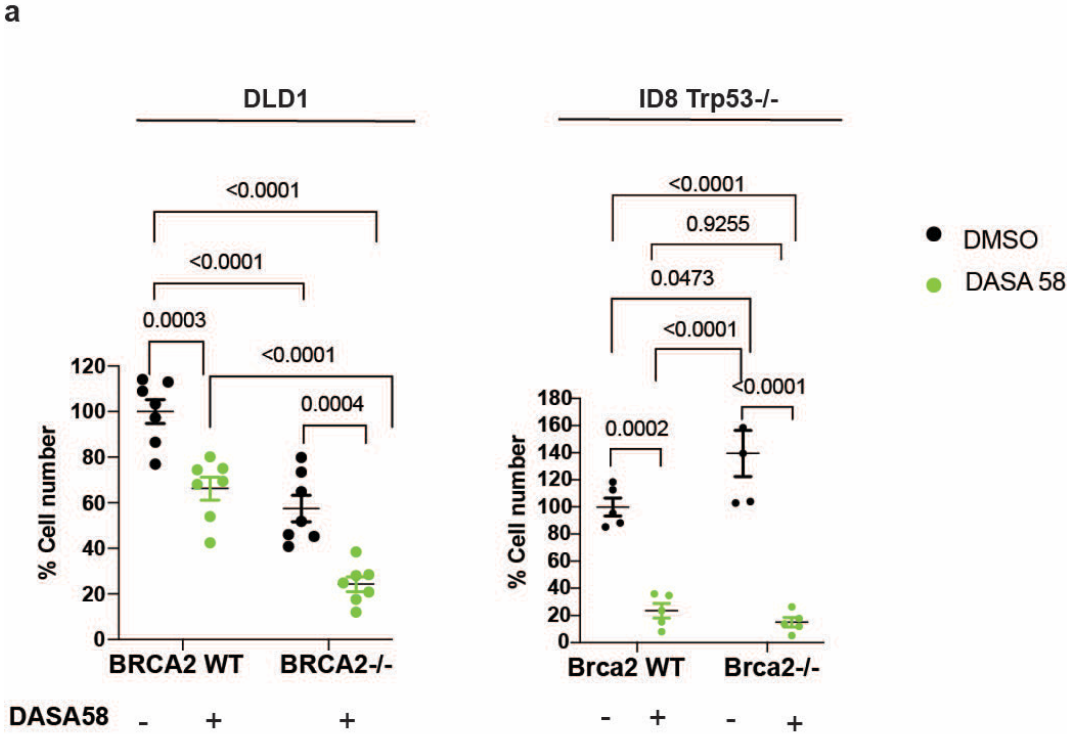

**Supplementary Fig.6. DASA58 exerts a cytostatic effect on both BRCA2 WT and BRCA2-deficient DLD1 tumor cells.** Cell number measured by Trypan blue assay in DLD1 BRCA2 WT and BRCA2<sup>-/-</sup> cells (left, n = 7) and ID8 Trp53<sup>-/-</sup> Brca2 WT and Brca2<sup>-/-</sup> cells (right, n=5) treated with DASA58 (50 μM) or DMSO. Data are presented as mean ± S.E.M. P values were determined by one-way ANOVA.

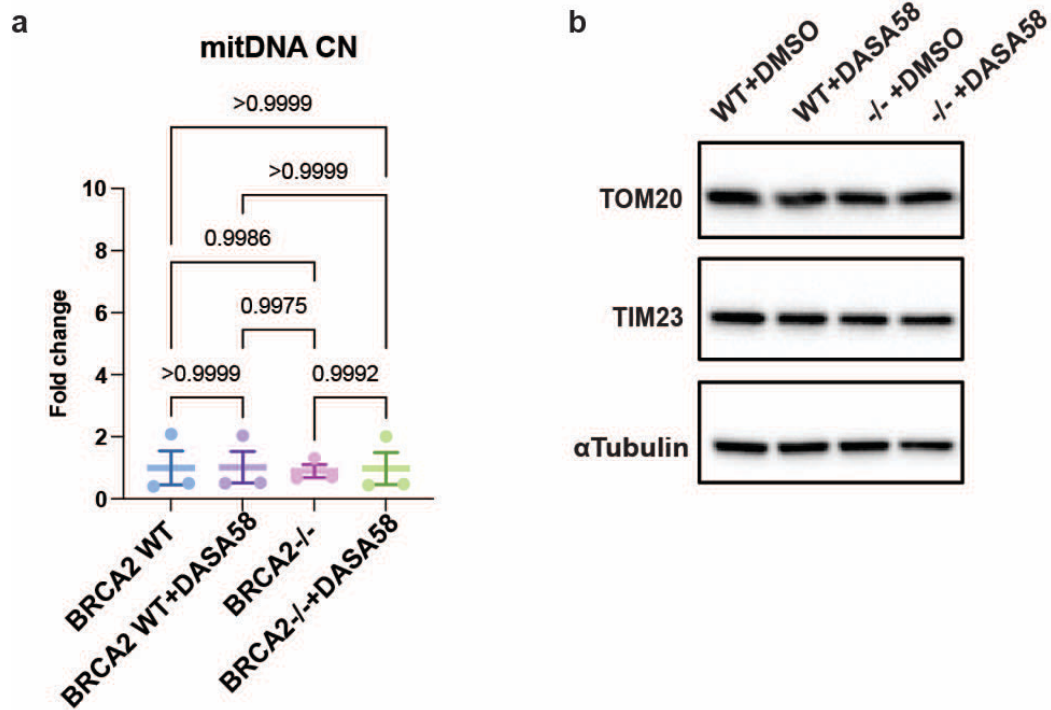

**Supplementary Fig.7. DASA58 and BRCA2 loss do not affect mitochondrial abundance.**
**a** Mitochondrial DNA copy number measured by qPCR in DLD1 BRCA2 WT and BRCA2<sup>-/-</sup> cells treated with or without DASA58 (n=3). **b** Immunoblot analysis of TOM20 and TIM23 in DLD1 BRCA2 WT and BRCA2<sup>-/-</sup> cells treated with or without DASA58. α-Tubulin was used as a loading control. A representative image of two independent cell lysates is shown.

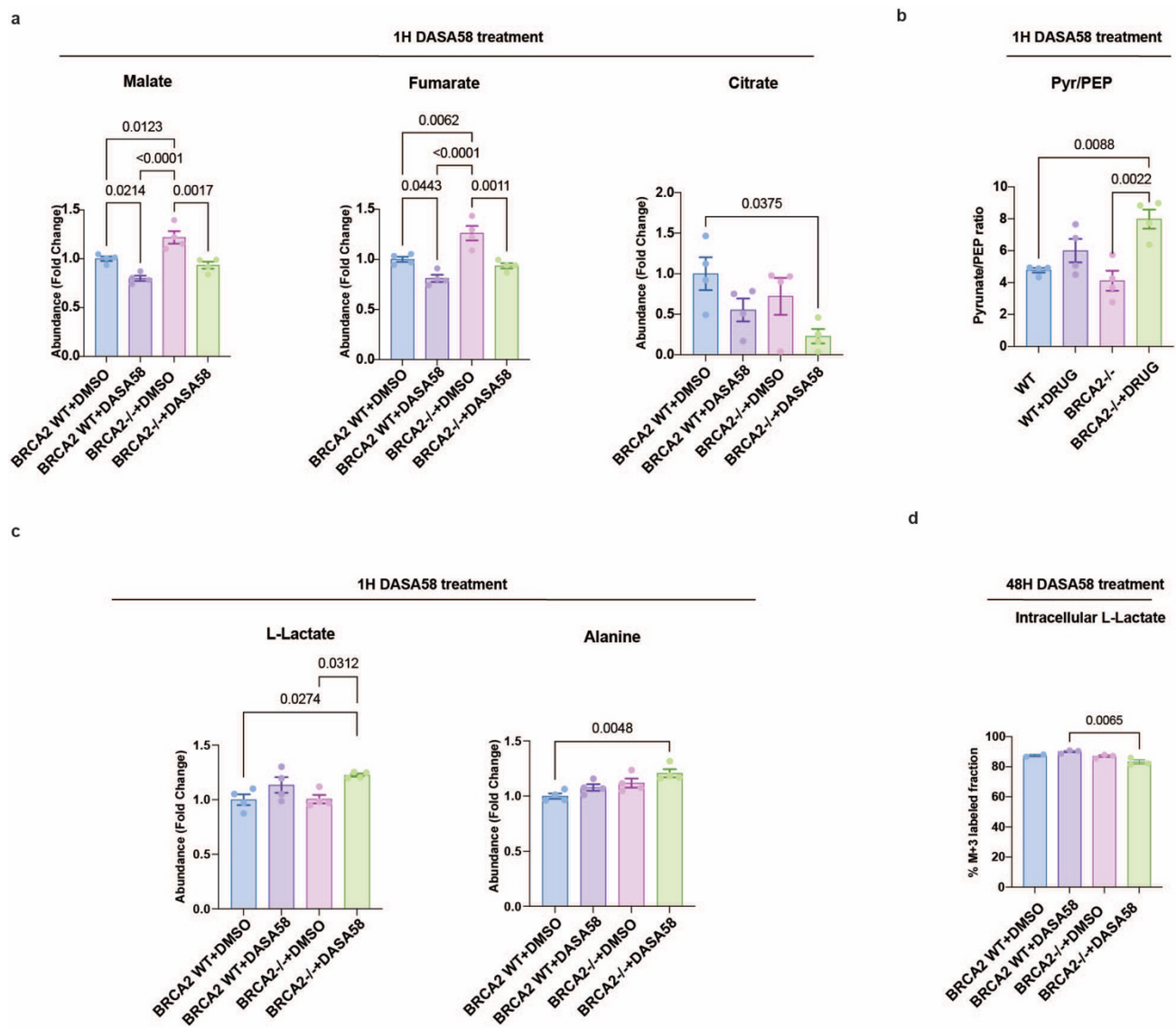

**Supplementary Fig.8. DASA58 treatment and BRCA2 loss alter the metabolite profile in DLD1.** Total abundance of malate, fumarate, and citrate, shown as fold change (**a**); pyruvate/phosphoenolpyruvate (PEP) ratio (**b**); and intracellular lactate and alanine levels (**c**) measured 1 h after DASA58 exposure (n=4). **d** M+3 labeled ($^{13}\text{C}$ incorporation) intracellular lactate levels measured 48 h after DASA58 exposure (30 minutes labelling) (WT n=2; WT+DASA58, BRCA2-/- and BRCA2-/-+DASA58, n=3). P values were determined by one-way ANOVA. Data are presented as mean ± S.E.M.

a

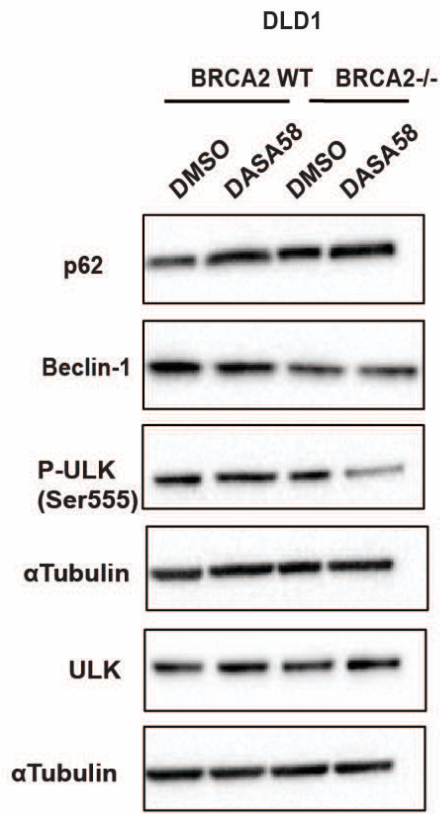

b

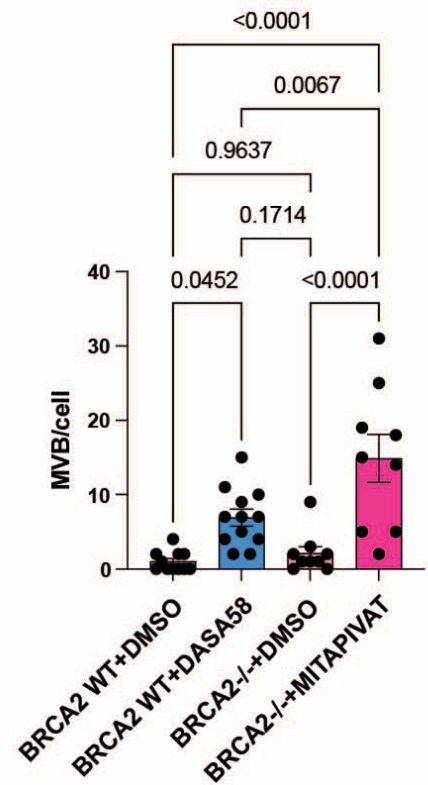

**Supplementary Fig.9. BRCA2 loss alters autophagy-associated markers, and DASA58 further enhances these effects.** **a** Representative Immunoblots showing p62, Beclin1, phospho-ULK, and total ULK levels in DLD1 BRCA2 WT and BRCA2<sup>-/-</sup> cells treated with DASA58 or vehicle control. A representative image of two independent cell lysates is shown. α-Tubulin served as a loading control. **b** Quantification of multivesicular body (MVB) number per cell from TEM images (n=9-12). *P* values were determined by one-way ANOVA. Data are presented as mean ± S.E.M.

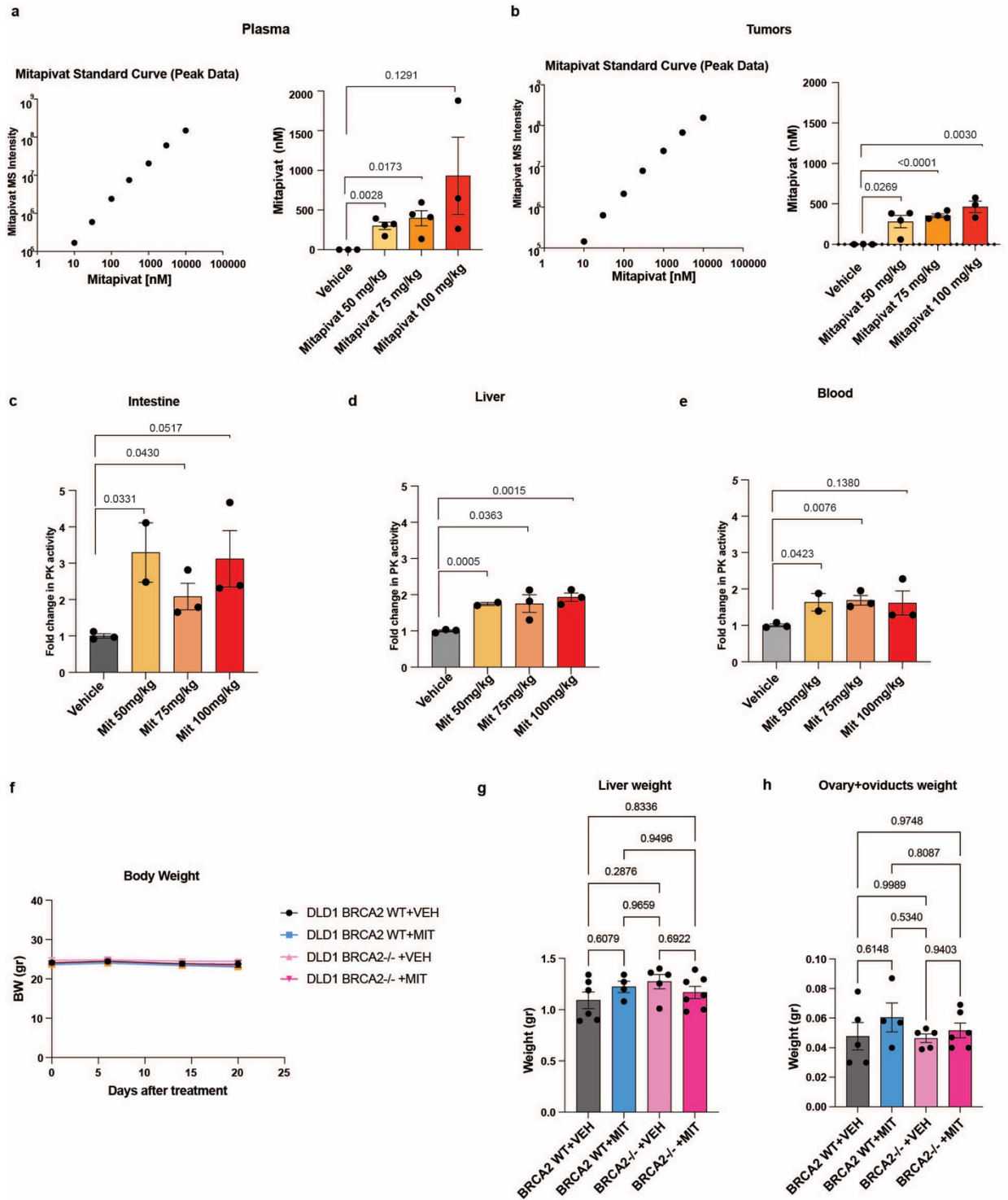

129

130 **Supplementary Fig.10. Mitapivat is bioavailable and does not cause detectable toxicity in**
131 **target organs.** Mitapivat levels detected in plasma (a) and tumors (b) by LC–MS at different
132 doses (50, 75, and 100 mg/kg) are shown, with the mitapivat standard curve in the left panel

(vehicle, mitapivat 50, 75 mg/kg n=4, 100mg/kg n=3). PK activity was measured in the intestine **(c)**, liver **(d)**, and blood **(e)** (vehicle, mitapivat 75, 100 mg/kg n=3, mitapivat 50 mg/kg n=2). **f** Body weight of nude mice bearing DLD1 tumors was recorded throughout the experiment (WT + vehicle, n = 8; WT + mitapivat, n = 8; BRCA2-/- + vehicle, n = 7; BRCA2-/- + mitapivat, n = 10). **g** Liver and **h** ovaries and oviducts weights were recorded at the end of the treatment (n=4-7). Data are presented as mean ± S.E.M. P values were determined by one-way ANOVA.

**Supplementary Table 1. Metabolomic dataset of DLD1 BRCA2 WT and BRCA2-/- cells treated with DASA58 for 1 hour.** Raw ion abundance values for a panel of 128 metabolites are shown.

**Supplementary Table 2. U-<sup>13</sup>C Glucose labeling in DLD1 BRCA WT and BRCA2-/- cells treated with DASA58 for 48h.** Raw ion abundance values and percent isotopologue abundance for lactate, malate, and citrate are shown are reported.
